# A cell fate decision map reveals abundant direct neurogenesis in the human developing neocortex

**DOI:** 10.1101/2022.02.01.478661

**Authors:** Laure Coquand, Anne-Sophie Macé, Sarah Farcy, Clarisse Brunet Avalos, Amandine Di Cicco, Marusa Lampic, Betina Bessières, Tania Attie-Bitach, Vincent Fraisier, Fabien Guimiot, Alexandre Baffet

**Affiliations:** Institut Curie, PSL Research University, CNRS UMR144, Paris, France; UMR 144-Cell and Tissue Imaging Facility (PICT-IBiSA), CNRS-Institut Curie, Paris, France; UF Embryofœtopathologie, Hopital Necker-enfants malades, Paris, France; UF de Fœtopathologie – Université de Paris et Inserm UMR1141, Hôpital Robert Debré, Paris, France; Institut national de la santé et de la recherche médicale (Inserm)

## Abstract

The human neocortex has undergone strong evolutionary expansion, largely due to an increased progenitor population, the basal radial glial (bRG) cells. These cells are responsible for the production of a diversity of cell types, but the successive cell fate decisions taken by individual progenitors remains unknown. Here, we developed a semi-automated live/fixed correlative imaging method to generate a map of bRG cell division modes in early fetal tissue and cerebral organoids. Through the analysis of over 1,000 dividing progenitors, we show that bRG cells undergo abundant symmetric amplifying divisions, followed by frequent direct neurogenic divisions, bypassing intermediate progenitors. These direct neurogenic divisions are more abundant in the upper part of the subventricular zone. We furthermore demonstrate asymmetric Notch activation in the self-renewing daughter cells, independently of basal fiber inheritance. Our results reveal a remarkable conservation of fate decisions in cerebral organoids, supporting their value as models of early human neurogenesis.

## Introduction

The human neocortex, composed of billions of neuronal and glial cells, is at the basis of higher cognitive functions (Cadwell et al., 2019). Its evolutionary size expansion is particularity important in the upper (supragranular) layer, leading to increased surface area and folding (Nowakowski et al., 2016). This is largely due to progenitor cells called basal radial glial cells (bRGs), also known as outer radial glial cells (Fietz et al., 2010; Hansen et al., 2010; Reillo et al., 2011; Smart et al., 2002). These cells are highly abundant in human - but rare in the lissencephalic mouse (Vaid et al., 2018; Wang et al., 2011) – and reside in the outer subventricular zone (OSVZ) where they contribute to the majority of supragranular neurons (Dehay et al., 2015).

bRG cells derive from apical (or ventricular) RG cells but have lost their connection to the ventricular surface through a process resembling an epithelial-mesenchymal transition (EMT) (**Figure 1A**) (LaMonica et al., 2013; Martínez-Martínez et al., 2016). A major feature of bRG cells is the presence of an elongated basal process along which newborn neurons migrate, though various morphologies have been reported including the presence of an apical process that does not reach the ventricle (Betizeau et al., 2013). bRG cells express various RG markers such as PAX6, Vimentin and SOX2, and undergo an unusual form of migration called mitotic somal translocation (MST) which occurs shortly before cytokinesis (Ostrem et al., 2014). Consistent with a steady increase of the bRG cell pool during development, live imaging experiments have documented their high proliferative potential (Betizeau et al., 2013; Fietz et al., 2010; Hansen et al., 2010). bRG cells are believed to increase the neurogenic output of the cortex, while providing extra tracks for radial migration and tangential dispersion of neurons (Fernández et al., 2016).

**Figure 1.**
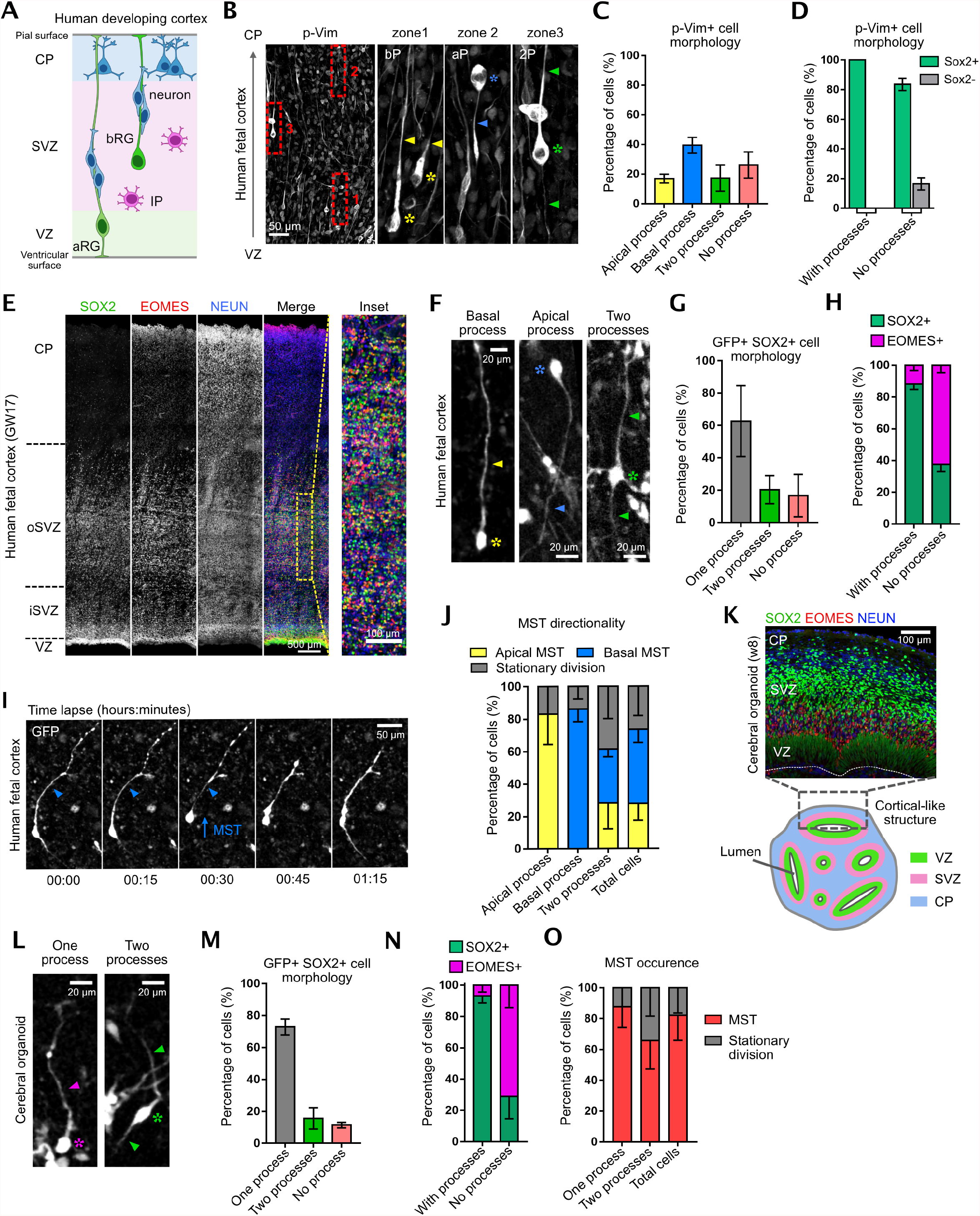
Morphological characterization of bRG cells in human cerebral organoids and fetal tissue. **A**. Schematic representation of human neocortex development. VZ: ventricular zone. SVZ: subventricular zone. CP: cortical plate. aRG: apical radial glial progenitor. bRG: basal radial glial progenitor. IP: Intermediate progenitor. **B**. Phospho-Vimentin immunostaining of human frontal cortex at GW18. Image is overexposed to visualize processes. Insets show three bRG cells with basal process (zone 1), apical process (zone 2) and two processes (zone 3). Asterix indicate soma and arrowheads indicate processes. **C**. Quantification of mitotic bRG cell morphologies in GW 14-18 frontal cortex. (N=3 brains, 338 cells). **D**. Percentage of p-VIM+ cells positive for SOX2, depending on morphology. (N=3 brains, 456 cells). **E**. SOX2, EOMES and NEUN immunostaining in human frontal cortex at GW17. **F**. Morphologies of GFP-expressing SOX2+ cells in human frontal cortex at GW17. **G**. Quantification of morphologies of GFP expressing SOX2+ cells in human frontal cortex at GW 14-17 (N=2 brains, 350 cells). **H**. Percentage of SOX2+ and EOMES+ progenitors, depending on morphology in human frontal cortex at GW 14-17 (N=2 brains, 204 cells). **I**. Live imaging of bRG cell performing MST in human fetal tissue. Arrowhead indicates basal process. **J**. Directionality of MST depending on bRG cell morphology in human frontal cortex at GW 14-18 (N=3 brains, 242 cells). **K**. (Top) SOX2, EOMES and NEUN immunostaining in week 8 cerebral organoid. (Bottom) Schematic representation of week 8-10 cerebral organoids. **L**. Morphologies of GFP expressing SOX2+ cells in cerebral organoids at weeks 7-10. **M**. Quantification of morphologies of GFP-expressing SOX2+ cells in cerebral organoids at weeks 7-8. (N=2 batches, 104 cells) **N**. Percentage of SOX2+ and EOMES+ progenitors, depending on morphology in cerebral organoids at weeks 8-10. (N=3 batches, 205 cells) **O**. Directionality of MST depending on bRG cell morphology cerebral organoids at weeks 8-9 (N=4 batches, 260 cells). Error bars indicate SD.

Genomic analyses have revealed the transcriptional profile of bRG cells as well as the cellular diversity in the human developing neocortex (Florio et al., 2015; Nowakowski et al., 2017; Pollen et al., 2015). They highlighted the conservation of cellular identities between fetal tissue and cerebral organoids, despite some degree of metabolic stress (Amiri et al., 2018; Bhaduri et al., 2020; Camp et al., 2015; Kanton et al., 2019; Pollen et al., 2019; Velasco et al., 2019). Such studies led to the identification of several human bRG-specific genes with important roles in bRG cell generation and amplification (Fiddes et al., 2018; Florio et al., 2015; Lui et al., 2014; Suzuki et al., 2018). How these factors affect bRG cell fate decisions *in vivo* remains however unclear.

Large scale genomic methods, or clonal methods such MADM or iTracer, are powerful tools to identify cellular diversity and global lineage relationships, but do not allow the probing of cell fate decisions taken by individual cells (Gao et al., 2014; He et al., 2021). Indeed, very different sequences of progenitor divisions could lead to the same final cellular output (Fischer and Morin, 2021). Identifying these progenitor cell fate decisions modes (i.e. the fate of their daughter cells) is critical to understand how neurogenesis is regulated across species, and affected in pathological contexts. Before gliogenic stages, bRG cells can, theoretically, undergo several division modes: symmetric proliferative (two RG daughters), symmetric self-consuming (two differentiating daughters) or asymmetric self-renewing divisions (one RG and one differentiating daughter). Moreover, differentiating divisions can lead to the production of a neuron (direct neurogenic division) or of an intermediate progenitor (IP, indirect neurogenic division). This IP can itself have different proliferative properties.

Here, we developed a method to quantitatively map human bRG cell division modes. Using a semi-automated live-fixed correlative imaging approach that enables the identification of the fate of dividing bRG daughters in space and time, we have established a map of cell fate decisions in human fetal tissue and cerebral organoids. We observe a remarkable similarity of division modes between the two tissues, and identify a major – although not exclusive – trajectory for bRG cells: symmetric amplifying divisions followed by self-consuming direct neurogenic divisions, independently of IP generation. Within asymmetrically dividing cells, we demonstrate that basal process inheritance does not correlate with self-renewal as it does in aRG cells, and is a consequence rather than a cause of bRG cell fate. We show that this is due to a difference in Notch signaling, possibly caused by the different microenvironments of these two cell types.

## Results

### Morphological identification of bRG cells

In order to identify human bRG cell division modes using live imaging methods, we first validated the identification of these cells based on morphological features. Human fetal pre-frontal cortex tissues from Gestational Week (GW) 14-18 were stained for phospho-Vimentin, which marks mitotic RG cells. Imaging within the SVZ revealed four different morphologies for these cells: unipolar with a single apical process (not reaching the VZ), unipolar with a basal process, bipolar with both an apical and a basal process, and cells with no visible process (**Figure 1B, 1C**). Mitotic bipolar bRG cells always had a major thick process and a minor thin process, which could be apical or basal (**Figure 1B**, zone 3). Overall, over 80% p-VIM+ cells displayed at least one process, and 60% a basal process. All process-harboring p-VIM+ cells were also SOX2+, while 20% of non-polarized p-VIM+ cells were negative for SOX2 (**Figure 1D**).

We then explored bRG cell morphology in non-mitotic cells marked with a cytoplasmic GFP. Fetal brains slices were infected with retroviruses (RV) and stained for SOX2 (RG cells), EOMES (IPs) and NEUN (Neurons) (**Figure 1E and S1A-S1B**). This analysis confirmed that over 80% of SOX2+/EOMES-/NEUN-cells displayed apical and/or basal processes, while 20% were non-polarized (**Figures 1F, 1G**). Moreover, the majority of process-harboring cells were SOX2+/EOMES-/NEUN-, and around 40% of non-polarized cells were SOX2+/EOMES-/NEUN-(**Figure 1H**). Therefore, human fetal bRG cells largely display elongated processes, though 20% are non-polarized.

We next performed live imaging of GFP-expressing cells in fetal slices, focusing on elongated bRG cells. Dividing cells had the same morphology as previously described in fixed samples (**Figure S1C**). The majority of process-harboring cells performed mitotic somal translocation (MST), though 25% performed stationary divisions (**Figure 1I, J; Video S1**). MST could occur in the apical direction or in the basal direction, depending on their shape. When bRG cells had two processes, MST occurred in the dominant (thick) process (**Figure 1J**).

Finally, we asked whether these morphological features were conserved in dorsal forebrain organoids. Week 8-10 organoids were infected with GFP-expressing retroviruses and stained for the cell fate marker SOX2, EOMES and NEUN, which revealed abundant SOX2+ bRG cells above the ventricular zone (**Figure 1K and S1D-S1E**). As in fetal tissue, the majority of SOX2+/EOMES-/NEUN-cells displayed one or two elongated processes, and 20% were non-polarized (**Figure 1L, M**). It was not possible to unambiguously identify whether processes were apical or basal, as bRG cells were often located between two lumens. The vast majority of process-harboring cells, and around 40% non-polarized cells, were SOX2+/EOMES-/NEUN-(**Figure 1N**). Live imaging confirmed these morphologies and indicated that the majority of bRG cells performed MST (**Figure 1O and S1F**). Therefore, the majority of human bRG cells can be identified in live samples based on their elongated morphology and ability to divide, which is conserved between fetal tissue and organoids.

### A semi-automated correlative microscopy method to identify cell fate decisions

We next developed a method to identify the fate acquired by daughter cells following progenitor cell division, in week 8-10 cerebral organoids. We established a correlative microscopy method consisting of live imaging GFP-expressing progenitors and, following fixation and immunostaining, assigning a fate to the live imaged cells (**Figure 2A**). The identification of corresponding cells between the live and fixed samples can be particularity challenging as the tissue is complex and multiple slices are imaged in parallel in up to 4 dishes (60-70 videos per acquisition). Moreover, slices rotate and even flip during the immunofluorescence process. We therefore developed a computer-assisted method to automate the localization of the videos in the immunostained samples (see methods). In brief, RV-infected tissue slices are live imaged for 48 hours and, at the end of the movie, 4X brightfield images of the slices containing positional information from each video are generated (**Figure 2B; Video S2**). Slices are then fixed, stained for the cell fate markers SOX2, EOMES and NEUN, and mosaic (tiled) images of the entire slices are acquired. Both live and fixed images are automatically segmented, paired, flipped and aligned. The position of each video is thereby obtained on the immunostained images, leading to the identification of matching cells between the live and fixed samples (**Figure 2B**). Using this method, bRG cells undergoing MST and dividing can be live imaged and the fate of the two daughter cells identified (**Figure 2C; Video S3**). Daughter cell fate was analyzed on average 30 hours after division. We noted that when a daughter cell differentiated (e.g. into an EOMES+ IP), it always retained some expression of the mother cell fate marker (SOX2) (**Figure 2C**). Expression of a novel fate marker was indeed very rapid, with EOMES or NEUN being detected in daughter cells that had divided 1-2 hours prior to the end of the movie (**Figure S2A**). Similarly, dividing IPs and migrating neurons could be live imaged and cell fate analyzed at the last timepoint (**Figure 2D, 2E; Videos S4, S5**). Therefore, this semi-automated correlative microscopy method allows the identification of cell fate markers in live imaged cerebral organoids, in a highly reproducible and quantitative manner.

**Figure 2.**
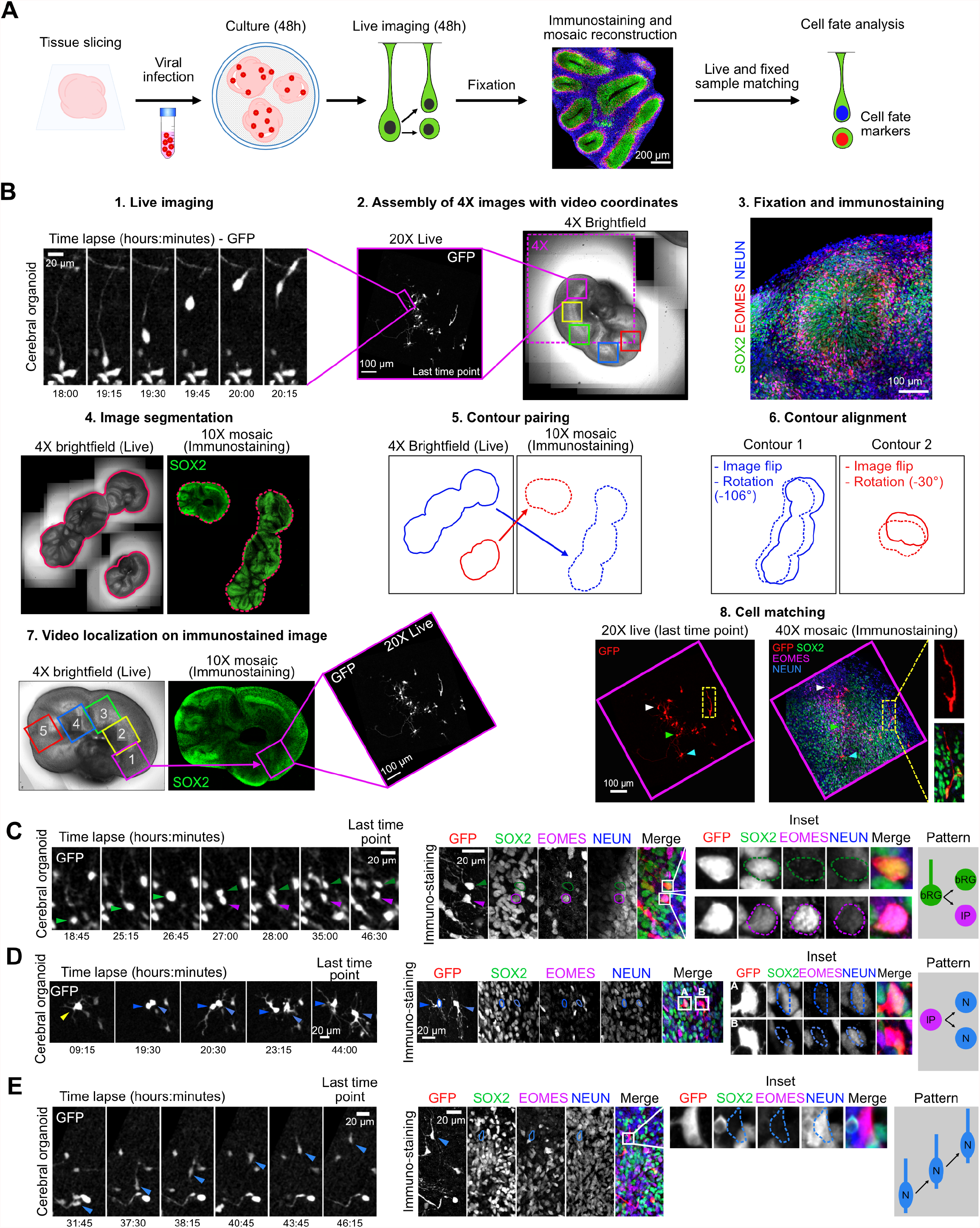
A semi-automated correlative imaging method to identify cell fate decisions in cerebral organoids. **A**. Schematic representation of correlative microscopy pipeline. **B**. Step-by-step protocol for semi-automated correlative microscopy. (1) bRG cells are live imaged for 48 hours. (2) 4X brightfield images containing the video coordinates are assembled. (3) Organoid slices are fixed, immunostained for SOX2, EOMES and NEUN and imaged. (4) Images are automatically segmented to outline slices from live and fixed samples. (5) Slice contours are automatically paired based on shape and area and (6) aligned (including a horizontal flip if needed). (7) Video fields of view are automatically annotated on the immunostaining images. (8) Regions of interest are re-imaged at higher resolution 40X and cells from live and fixed samples are manually matched. **C**. Live/fixed correlative analysis of a dividing bRG cell generating a self-renewing bRG daughter and a differentiating IP daughter. **D**. Live/fixed correlative analysis of a dividing IP cell generating two neuronal daughters. **E**. Live/fixed correlative analysis of a migrating neuron.

### A map of cell fate decisions in cerebral organoids

To generate a map of progenitor division modes, we analyzed 444 dividing bRG cells, in week 8-10 cerebral organoids, prior to the start of gliogenesis (Pollen et al., 2019; Velasco et al., 2019) (**Figures 3A, 3B, 3C ; Videos S6, S7, S8**). We first quantified the fraction of proliferative divisions (leading to two SOX2+ cells) versus neurogenic divisions (leading to at least one differentiating cell, EOMES+ or NEUN+). This analysis revealed a high rate of bRG cell amplification at this stage, with over 60% of proliferative divisions (**Figures 3D, 3E**). Within neurogenic divisions, different patterns could be observed. bRG cells performed symmetric self-consuming divisions, leading to two differentiating cells, or asymmetric self-renewing divisions, leading to one bRG cell and one differentiating cell. Self-consuming divisions were quite frequent (close to 40% of neurogenic divisions) and therefore represent an important mode of neuronal generation by bRG cells (**Figures 3D, 3F**). In both types of neurogenic division (asymmetric or symmetric), bRG cells could divide directly into neurons or indirectly, via the generation of IPs. Strikingly, we observed abundant direct neurogenesis by bRG cells (60% of neurogenic divisions), indicating that generation of IPs is not a systematic differentiation trajectory in these cells (**Figures 3D, 3G**). Notably, we never observed asymmetrically dividing bRG cells generating one IP and one neuron. We next performed the correlative microscopy analysis on dividing IPs. These cells were observed to perform over 60% of symmetric proliferative divisions, indicating a strong amplification potential (**Figure S2B**). Overall, this analysis indicates that, in week 8-10 cerebral organoids, bRG cells are highly proliferative and undergo important self-amplification. Upon differentiation, they undergo different neurogenic routes, with frequent self-consuming terminal divisions, as well as abundant direct neurogenesis.

**Figure 3.**
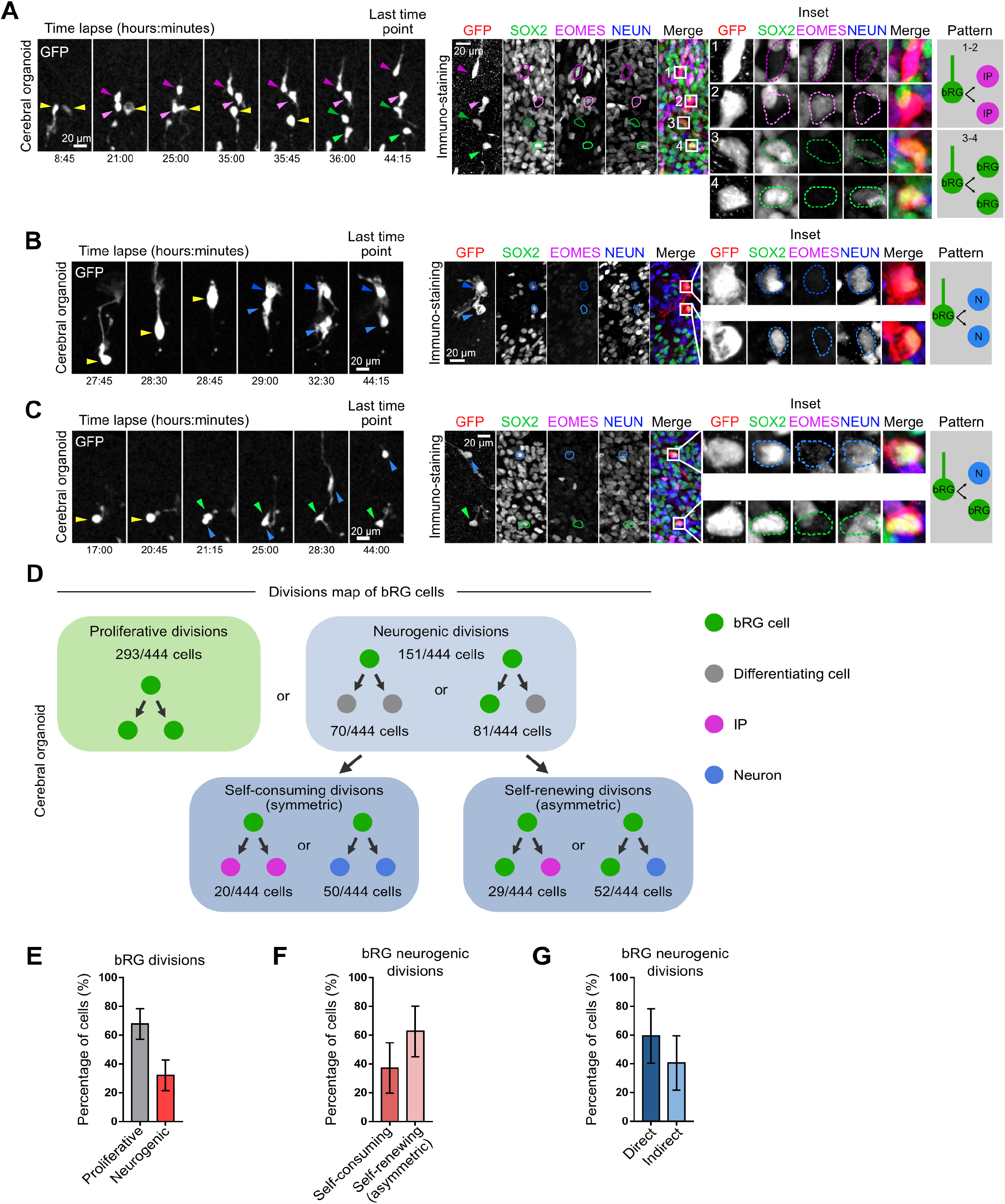
A map of cell fate decisions in human cerebral organoids. **A**. (Top) Live/fixed correlative analysis of a dividing bRG cell generating two IP daughters. (Bottom). Live/fixed correlative analysis of a dividing bRG cell generating two bRG daughters.**B**. Live/fixed correlative analysis of a dividing bRG cell generating two neuronal daughters. **C**. Live/fixed correlative analysis of a dividing bRG cell generating a bRG daughter and a neuronal daughter. **D**. Summary of all division patterns identified in bRG cells within week 8-10 cerebral organoids (N=444 bRG cells from 3 batches). **E**. Quantification of the percentage of proliferative versus neurogenic divisions of bRG cells in week 8-10 cerebral organoids. (N=5 batches, 444 cells). **F**. Quantification of the percentage of self-consuming versus asymmetric self-renewing divisions of bRG cells in week 8-10 cerebral organoids (N=5 batches, 151 cells).**G**. Quantification of the percentage of direct versus indirect neurogenic divisions of bRG cells in week 8-10 cerebral organoids (N=5 batches, 151 cells). Error bars indicate SD.

### A map of cell fate decisions in human fetal tissue

We next adapted this correlative microscopy method to human frontal cortex samples at GW 14-17. These stages were selected to match week 8-10 cerebral organoids, based on transcriptomics and because both are mostly pre-gliogenic (Pollen et al., 2019). While slices were substantially larger, the macro proved to be very efficient at automatically identifying and aligning corresponding regions between the live and fixed datasets (**Figures 4A, 4B**). We analyzed the division modes of 522 human fetal bRG cells, following 48-hour live imaging (**Figures 4C, 4D; Videos S9, S10**). We confirmed rapid expression of differentiation markers following cell division (**Figure S3A**). As in cerebral organoids, the majority of bRG cells performed symmetric proliferative division (70%), generating two SOX2+ daughters (**Figures 4E, 4F**). Within the neurogenic divisions, we again observed abundant symmetric self-consuming divisions (40% of all neurogenic divisions) (**Figures 4E, 4G**). We confirmed that direct neurogenic divisions are a major bRG cell division mode (over 40% of all neurogenic divisions) (**Figures 4E, 4H**). Finally, we analyzed the division modes of human IPs in fetal samples. These cells also demonstrated a high proliferative potential, with 85.7% symmetric amplifying divisions (**Figure S3B**). Therefore, division patterns in week 14-17 fetal cortex closely match those of week 8-10 cerebral organoids, with bRG cells showing strong amplification potential and, upon differentiation, frequent direct neurogenic divisions and self-consuming divisions.

**Figure 4.**
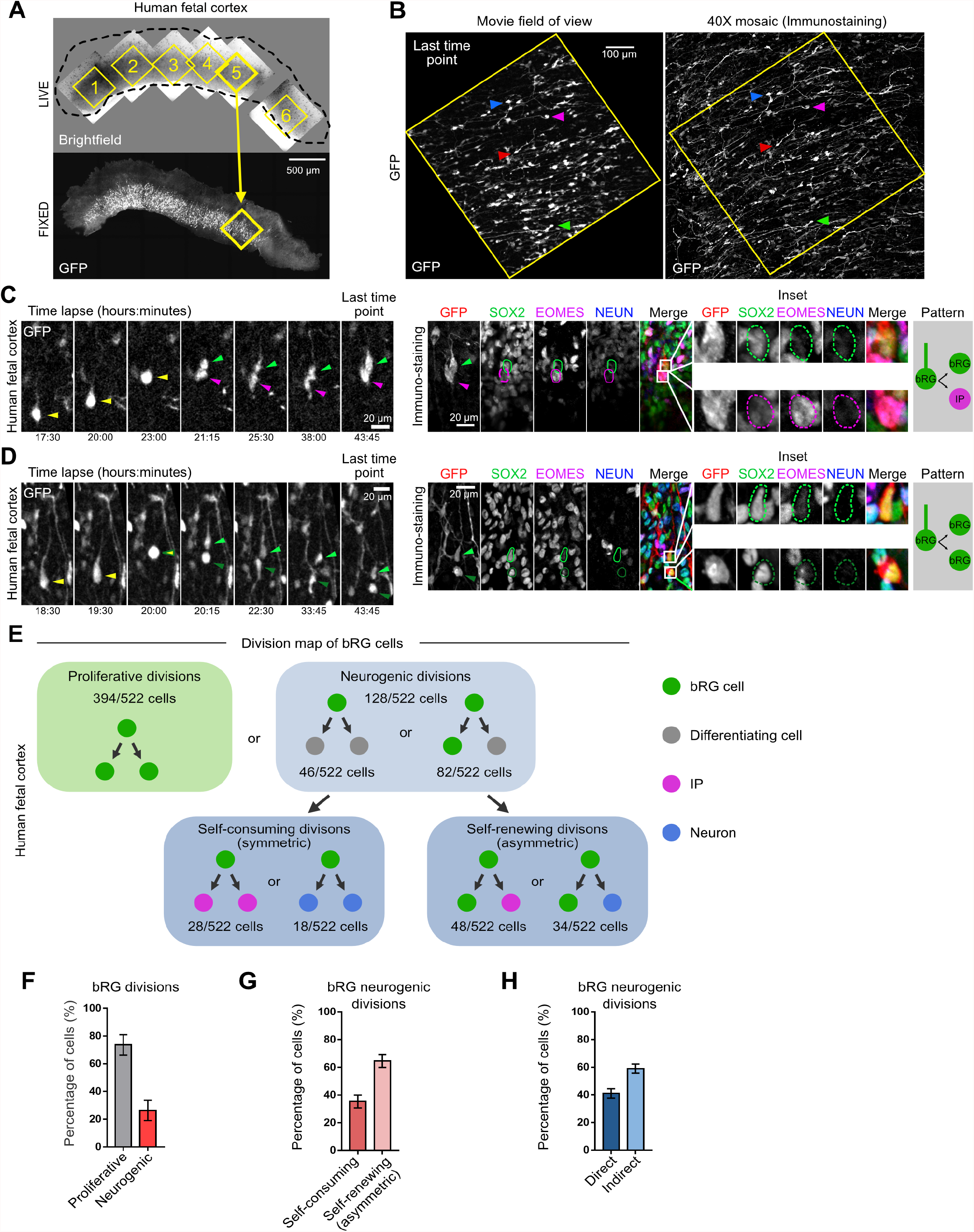
A map of cell fate decisions in fetal human frontal cortex. **A**. Automated pairing of live and fixed samples and annotation of the video fields of view on the immunostained fixed samples. **B**. GFP+ cell matching between the live images and the fixed images. Arrowheads indicate equivalent cells. **C**. Live/fixed correlative analysis of a dividing bRG cell generating a bRG daughter and an IP daughter. **D**. Live/fixed correlative analysis of a dividing bRG cell generating two bRG daughters. **E**. Summary of all division patterns identified in bRG cells within GW 14-17 human frontal cortex (N=522 bRG cells from 2 fetal tissues). **F**. Quantification of the percentage of proliferative versus neurogenic divisions of bRG cells in GW 14-17 human frontal cortex (N=522 bRG cells from 2 fetal tissues). **G**. Quantification of the percentage of self-consuming versus asymmetric self-renewing divisions of bRG cells in GW 14-17 human frontal cortex (N=128 bRG cells from 2 fetal tissues). **H**. Quantification of the percentage of direct versus indirect neurogenic divisions of bRG cells in GW 14-17 human frontal cortex (N=128 bRG cells from 2 fetal tissues). Error bars indicate SD.

### Increased direct neurogenesis in the basal part of the human fetal OSVZ

The human OSVZ is extremely large (approximately 3 mm at GW17) and bRG cells may therefore be exposed to different microenvironments depending on their position, which may influence their division modes. Moreover, bRG cells progressively migrate through the SVZ and have a different history depending on their position. We therefore explored whether bRG division modes vary along the apico-basal axis in the human fetal brain. To test this, we adapted the above-described macro to automatically record the position of each dividing bRG cell within the tissue. Distance to the apical surface was measured at the time of cytokinesis. We then plotted the different division modes depending on bRG cell position within the tissue. The position of bRG cells along the apico-basal axis did not influence the rate of symmetric proliferative versus neurogenic division (**Figure 5A, 5B, 5C**). Similarly, the rate of symmetric self-consuming versus asymmetric self-renewing divisions was not apparently different (**Figure 5D, 5E, 5F**). However, we observed a significant difference in the rates of direct versus indirect neurogenesis. Indeed, indirect neurogenic divisions (EOMES+ cells) occurred on average 736 !m from the apical surface, while direct neurogenic divisions (NEUN+ cells) occurred much more basally, 1,114 !m from the apical surface (**Figure 5G, 5H, 5I**). Therefore, at this developmental stage, bRG self-amplification is abundant and does not vary along the apico-basal axis. Dividing bRG cells however undergo more direct neurogenic divisions when located in the basal part of the SVZ.

**Figure 5.**
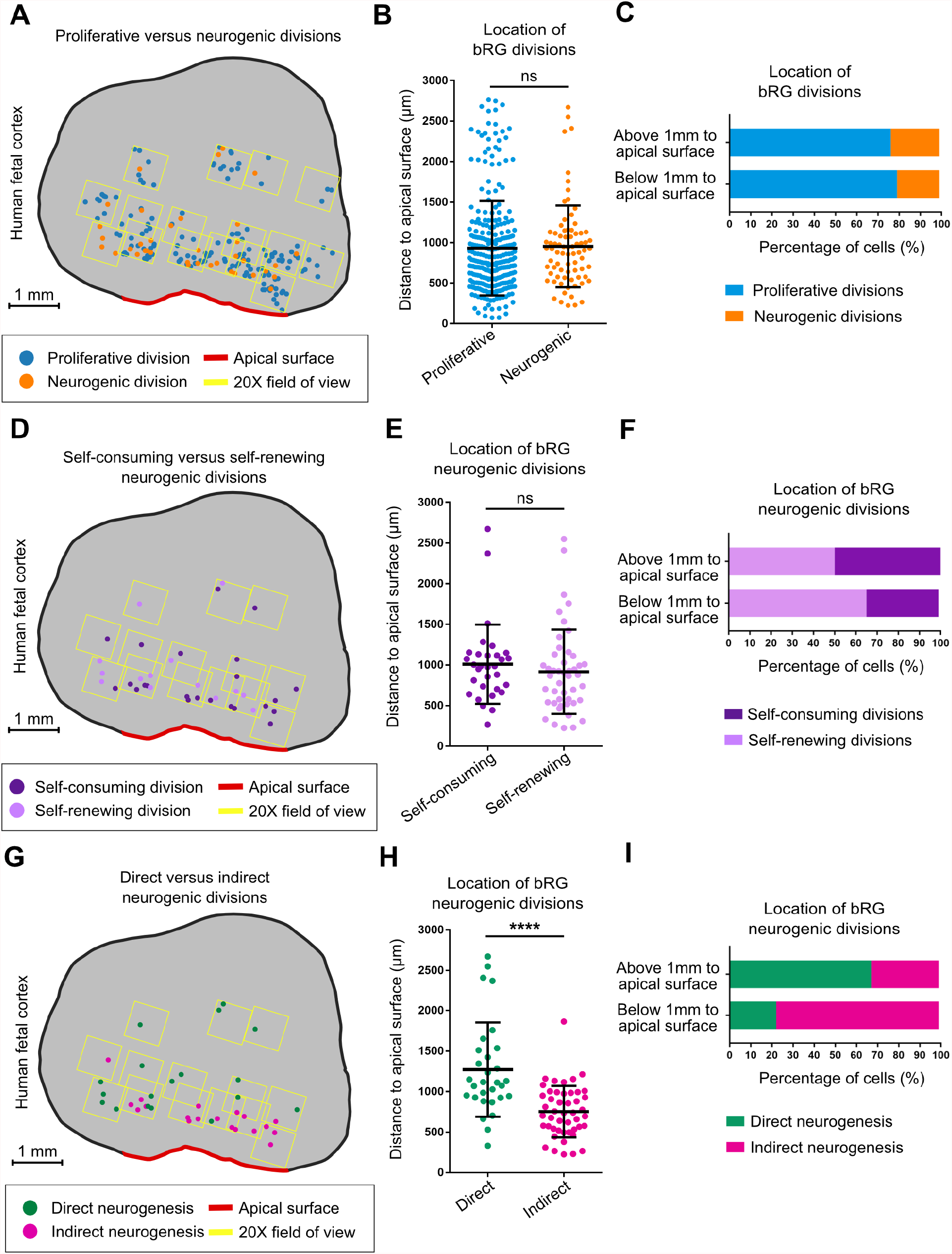
Spatial distribution of division modes in human fetal cortex. **A**. Spatial distribution of proliferative versus neurogenic divisions in GW17 human frontal cortex. **B**. Quantification of proliferative versus neurogenic divisions (N=355 cells from 1 fetal sample) **C**. Percentage of proliferative versus neurogenic divisions, below or above 1 mm to the apical surface (N=355 cells from 1 fetal sample). **D**. Spatial distribution of self-consuming versus asymmetric self-renewing divisions in GW17 human frontal cortex. **E**. Quantification of self-consuming versus asymmetric self-renewing divisions (N=75 cells from 1 fetal sample).**F**. Percentage of self-consuming versus asymmetric self-renewing divisions, below or above 1 mm from the apical surface (N=75 cells from 1 fetal sample). **G**. Spatial distribution of direct versus indirect neurogenic divisions in GW17 human frontal cortex. **H**. Quantification of direct versus indirect neurogenic divisions (N=75 cells from 1 fetal sample). **I**. Percentage of direct versus indirect neurogenic divisions, below or above 1 mm from the apical surface (N=75 cells from 1 fetal sample).

### Basal process inheritance does not predict bRG fate upon asymmetric division

The mechanism of bRG cell asymmetric division remains unknown. In mouse aRG cells, growing evidence support the role of basal process inheritance in stem cell fate maintenance (Alexandre et al., 2010; Peyre and Morin, 2012; Shitamukai et al., 2011). We therefore used our correlative imaging method to test whether process inheritance correlates with bRG fate maintenance upon asymmetric division of human bRG cells. We first live imaged 79 asymmetrically-dividing bRG cells (one bRG daughter – one differentiating daughter) within week 8-10 cerebral organoids, and analyzed daughter cell fate depending on process inheritance (**Figures 6A, 6B; Videos S11, S12**). In half of these cells, process-inheriting daughters maintained a bRG fate but in the other half, process-inheriting daughters differentiated (**Figures 6C, 6D**). This was the case whether the asymmetric divisions generated an IP or directly a neuron. These results suggest no role for process inheritance in bRG fate upon asymmetric cell division in cerebral organoids. We next performed a similar analysis in GW 14-17 human fetal brain tissue. We analyzed 82 asymmetrically dividing bRG cells and again found no correlation between basal process inheritance and bRG cell fate (**Figures 6E, 6F; Videos S13, S14**) 52.4% of basal process-inheriting daughters remained bRG cells, and 47.6% differentiated (**Figures 6G, 6H**). We did not observe any effect of the apical process on cell fate either (not shown). In support of these results, SOX2+ daughter cells that did not inherit a process could be observed to regrow a novel basal process after division (**Figure S4A; Video S15**). Therefore, in human bRG cells, the basal process appears to be a consequence, rather than a cause, of bRG cell fate upon asymmetric division. Its presence during interphase may however participate in long-term bRG fate maintenance.

**Figure 6.**
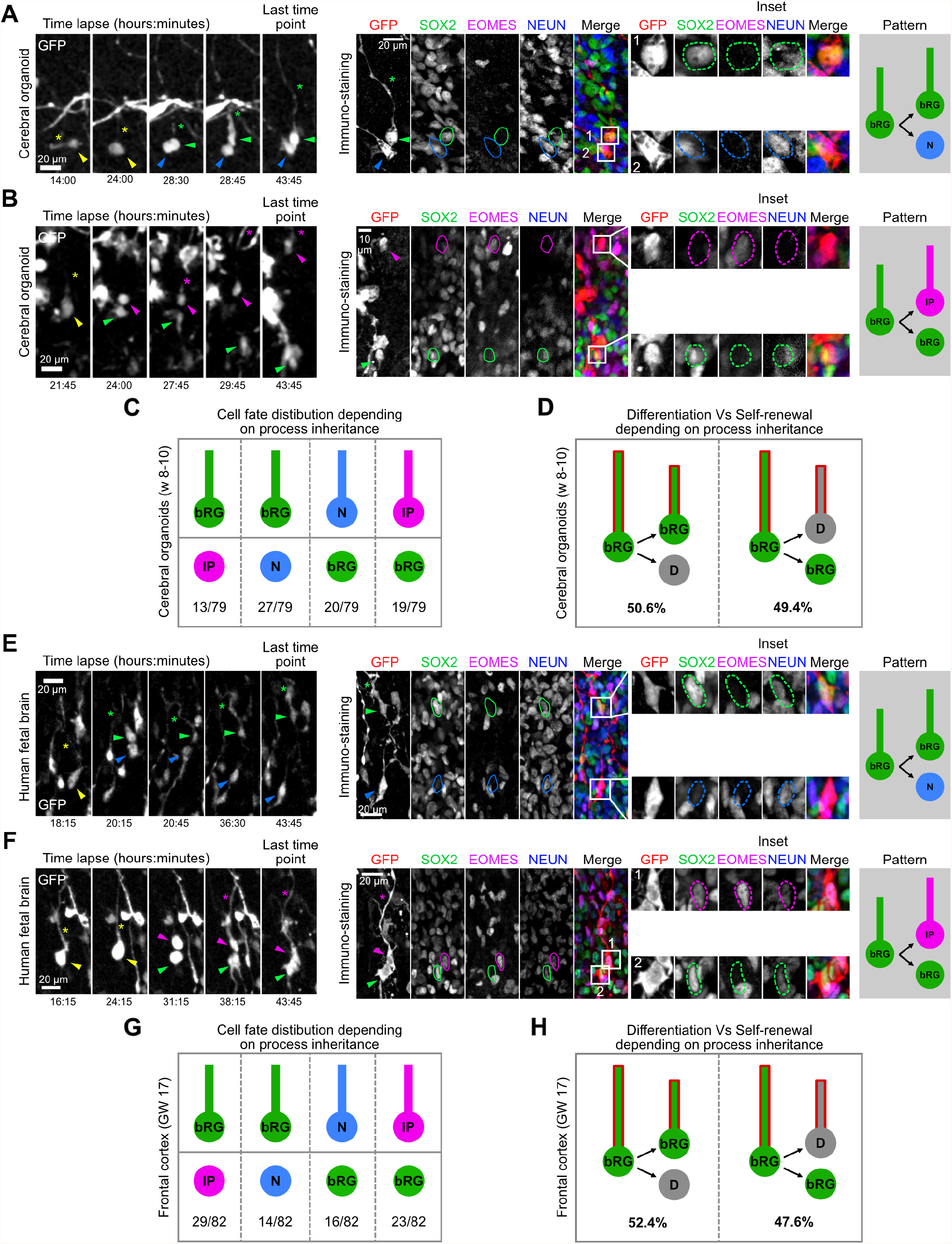
Basal process inheritance does not predict bRG fate upon asymmetric division. **A**. Live/fixed correlative analysis of basal process inheritance in a dividing bRG cell generating a process-inheriting bRG daughter and neuron, within a cerebral organoid. **B**. Live/fixed correlative analysis of basal process inheritance in a dividing bRG cell generating a process-inheriting IP daughter and a bRG daughter, within a cerebral organoid. **C**. Distribution of cell fates depending on process inheritance upon asymmetric cell division in week 8-10 cerebral organoids (N=79 asymmetrically dividing cells from 5 experiments). **D**. Percentage of self-renewing versus differentiating daughter cells upon asymmetric division, depending on process inheritance in week 8-10 cerebral organoids (N=79 asymmetrically dividing cells from 5 experiments). **E**. Live/fixed correlative analysis of basal process inheritance in a dividing bRG cell generating a process-inheriting bRG daughter and a neuron, within fetal frontal cortex. **F**. Live/fixed correlative analysis of basal process inheritance in a dividing bRG cell generating a process-inheriting IP daughter and a bRG daughter, within fetal frontal cortex. **G**. Distribution of cell fates depending on process inheritance upon asymmetric cell division in GW 14-17 human frontal cortex (N=82 asymmetrically dividing cells from 2 experiments). **H**. Percentage of self-renewing versus differentiating daughter cells upon asymmetric division, depending on process inheritance in GW 14-17 human frontal cortex (N=82 asymmetrically dividing cells from 2 experiments).

### Notch signaling is active in bRG daughters, not in process-inheriting cells

We next addressed why basal process inheritance correlates with stem cell fate in mouse aRG cells but not in human bRG cells. In aRG cells, it was proposed that the basal process acts as an antenna for the reception of Notch signaling from the surrounding cells, in particular neurons (Peyre and Morin, 2012; Shitamukai et al., 2011). We therefore investigated Notch signaling in bRG daughter cells, depending on process inheritance. As a readout, we analyzed expression of its downstream target HES1. In cerebral organoids, HES1 was strongly expressed in the VZ where aRG cells are highly abundant and in a sparse manner in the SVZ, reflecting the SOX2+ bRG cell distribution (**Figure 7A**). Week 8-11 organoid slices were live imaged for 48 hours, stained for HES1, EOMES and NEUN, and processed through the correlative imaging protocol. Cell fate was determined based on EOMES and NEUN expression, double-negative cells being identified as bRG cells. Out of 276 bRG cell, 186 performed symmetric proliferative divisions, 53 asymmetric divisions, and 37 symmetric self-consuming divisions (**Figure 7B**). Consistent with its oscillatory behavior in RG cells (Ochi et al., 2020), HES1 was only detected in a subset of bRG cells, whether these cells were generated following symmetric or asymmetric divisions (**Figures 7C, 7D; Video S16**). As expected, HES1 was never detected in differentiating cells (n=90 cells) (**Figure 7D**). In total, out of 276 live imaged bRG cells, we identified 16 cells that divided asymmetrically, with detectable HES1 expression in daughter cell (**Figure 7D**). HES1 was always detected in the non-differentiating daughter (EOMES-and NEUN-), supporting preferential Notch signaling in the self-renewing bRG daughter upon asymmetric division (**Figure 7D**). We found no correlation between HES1 expression and process inheritance: 8 HES1-expressing cells inherited the basal process and 8 did not (**Figure 7E, 7F; Video S17**). These data further support that process inheritance does not correlate with bRG cell fate, and that the basal process is not involved in differential Notch signaling upon asymmetric division in bRG cells, as it is believed to in aRG cells. We propose that this is due to the different microenvironments in which the soma of aRG and bRG cells localize (**Figure 7G**).

**Figure 7.**
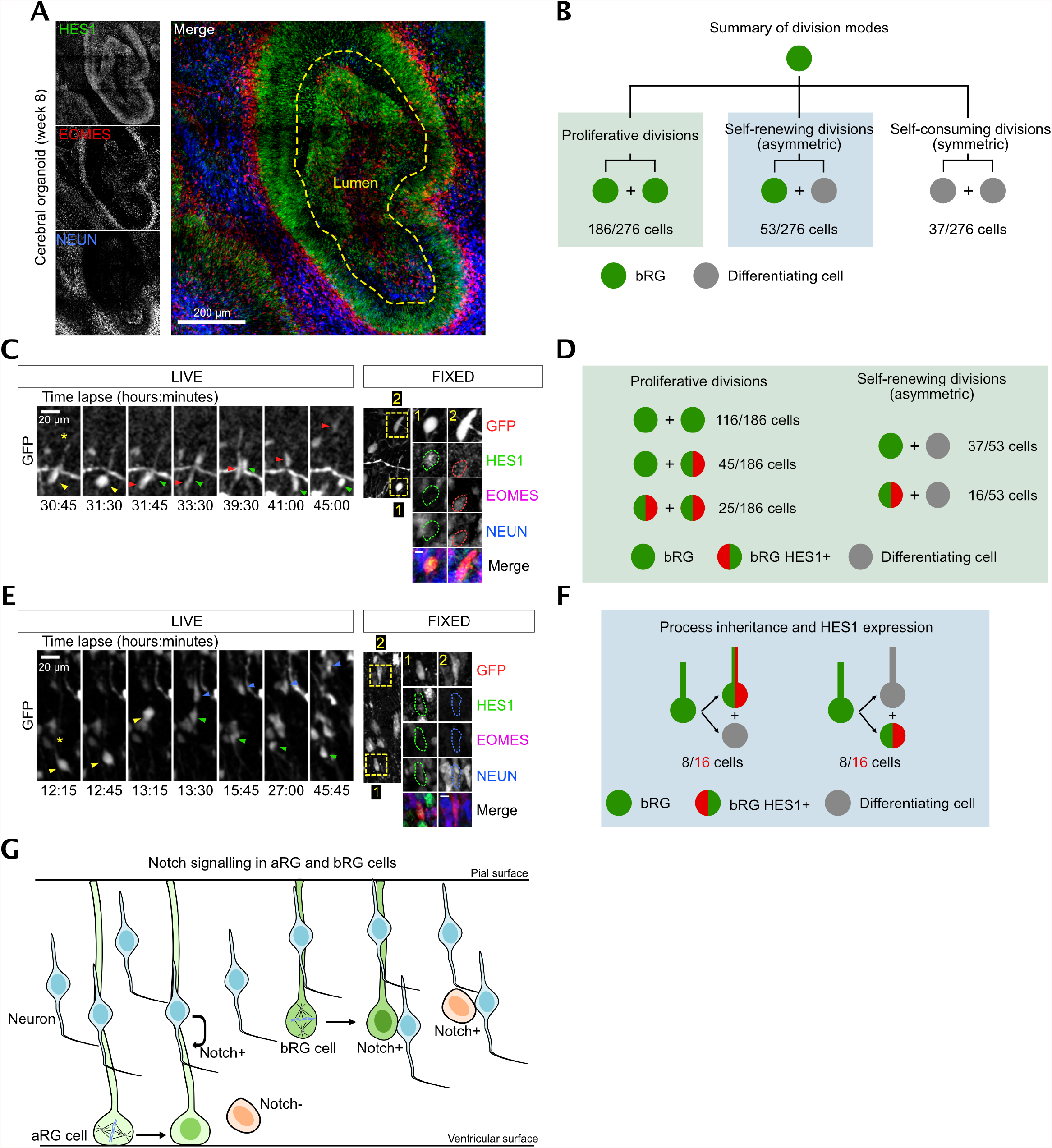
HES1 is preferentially expressed in bRG daughters, irrespective of process inheritance. **A**. HES1, EOMES and NEUN immunostaining in human cerebral organoid at week 8. **B**. Distribution of division modes identified in bRG cells within week 8-11 cerebral organoids. bRG daughter (EOMES- and NEUN-), differentiating daughter (EOMES+ or NEUN+) (N=276 bRG cells from 3 batches of organoids). **C**. Live/fixed correlative analysis of an asymmetrically dividing bRG cell revealing HES1 expression specifically in self-renewing daughter (EOMES- and NEUN-). **D**. Summary of HES1 expression in daughter cells depending on division modes (N= 239 cells from 3 batches of organoids). **E**. Live/fixed correlative analysis in asymmetrically dividing bRG cells revealing lack of correlation between HES1 expression and basal process inheritance. **F**. Summary of HES1 expression depending on process inheritance in asymmetrically dividing bRG cells, within week 8-11 cerebral organoids (N= 16 cells from 3 batches of organoids). **G**. Model for Notch activation in aRG and bRG cells. In aRG cells, basal process inheritance leads to differential Notch signaling between daughters. In bRG cells, the soma of both daughters is located in a neuron-rich region, impairing differential Notch signaling by the basal process.

## Discussion

bRG cells are key actors in the evolutionary expansion of the human brain, but the sequence of events leading to their massive neuronal output is unknown. Using live/fixed correlative imaging, we provide a map of their division modes at early – pre-gliogenic – stages. Identifying the precise cell fate decisions that lead to given neuronal outputs is critical to understand the diversity of differentiation trajectories taken by bRG cells. In mouse, aRG cells undergo a precise switch in division modes at E12.5, from mostly symmetric amplifying divisions to mostly asymmetric divisions generating one self-renewing aRG cell and one IP that will divide once to generate two neurons (Haubensak et al., 2004; Miyata et al., 2004; Noctor et al., 2004). Here we show that at GW 14-17 of human neocortex development, multiple bRG cell division modes co-exist, pointing to a more complex regulation of neurogenesis. At these stages, bRG amplification through symmetric cell divisions is dominant, in agreement with the increase of the bRG cell pool during early development (Fietz et al., 2010; Hansen et al., 2010). This result confirms that the high proliferative potential of human bRG cells is an important criterion to define them, in addition to morphology, localization and fate. bRG amplification may nevertheless substantially vary between species (Wang et al., 2011).

Our results indicate that a major trajectory for bRG cells consists of symmetric amplifying divisions, followed by self-consuming divisions that generate neurons directly, independently of IPs. These results are consistent with the lower abundance of EOMES+ cells in the human OSVZ, as compared to the ISVZ (Fietz et al., 2010). This represents another major difference with mouse aRG cells that largely rely on IPs to amplify the neurogenic output. Evolution of cortical neurogenesis in amniotes is regulated by the balance between direct and indirect neurogenesis (Cárdenas et al., 2018). aRG cells in sauropsids undergo direct neurogenesis, while mammals largely rely on indirect divisions in the evolutionary more recent neocortex, a process associated with size expansion and regulated by Robo signaling levels (Cárdenas et al., 2018). We show that this rule does however not apply to bRG cells, in which direct neurogenesis is common. aRG cells rely on IPs to amplify their neurogenic output, as their own amplification is limited by spatial constraints. They must indeed divide at the ventricular surface to precisely segregate their apical junctions between daughters and maintain a proper neuroepithelial structure. Interkinetic Nuclear Migration (INM) leads to the formation of a pseudostratified epithelium that allows an increase in the aRG cell pool, but their amplification still reaches a physical limit (Baffet et al., 2015; Hu et al., 2013; Lee and Norden, 2013). bRG cells on the other hand are not subject to this physical limitation and can amplify their own pool both radially and tangentially, and thus IPs are less relied upon to increase the neurogenic output. Whether direct and indirect divisions ultimately lead to the formation of different neuronal subtypes, as observed in aRG cells, remains to be tested (Cárdenas et al., 2018).

Cerebral organoids have emerged as a powerful system to investigate human brain development (Lancaster et al., 2013; Paşca et al., 2015; Qian et al., 2016). To what degree they faithfully recapitulate fetal neurogenesis is however important to monitor. Genomics studies have highlighted the similarity of transcriptional profiles, though substantial metabolic stress has been reported in organoids (Amiri et al., 2018; Bhaduri et al., 2020; Camp et al., 2015; Kanton et al., 2019; Pollen et al., 2019; Velasco et al., 2019). Here, we report a high similarity of bRG cell division modes between week 8-10 organoids and gestational week 14-17 fetal tissue. Importantly, an advantage of imaging approaches such as ours is that the necrotic core (from where most stress likely originates) can be avoided, focusing on the cortical-like lobes at the periphery of the organoids. These cortical-like structures are however much thinner than in the fetal brain, limiting the ability to probe how bRG cell position impacts their division modes, as performed here in fetal tissue. An open question is whether cell fate decisions remain highly similar at later stages of development.

The molecular mechanism regulating asymmetrical division in RG cells has been a matter of controversy. In aRG cells, increasing evidence support a role for the basal process in cell fate, which correlates with Notch activation and self-renewal (Alexandre et al., 2010; Peyre and Morin, 2012; Shitamukai et al., 2011). We however do not observe such a correlation in human bRG cells where Notch signaling is activated in the self-renewing daughter irrespective of basal process inheritance. aRG somas are located in the ventricular zone and their basal process extends through the cortex, contacting neurons from which Notch-Delta signaling can be activated. bRG somas on the other hand are located in the SVZ and both their daughter cells are in close proximity to neurons (**Figure 7G**). Therefore, due to the bRG cell microenvironment, it is consistent that their basal process does not confer differential Notch signaling. Other factors, such as centriole age, mitochondrial dynamics, mitotic spindle positioning or Sara endosomes are promising candidates (Iwata et al., 2020; Kressmann et al., 2015; Wang et al., 2009).

Descriptions of clonal relationships are a powerful means to understand cellular diversity. Key to this is the identification of the cell fate decision branch points along lineages.

The semi-automated correlative microscopy method enables us to quantitatively measure progenitor cell division modes in human cortical tissue. This will allow to probe neuronal subtype generation or the switch to gliogenesis, through time and space, across species, and in pathological contexts.

## Material and methods

### Ethics statement

Human fetal tissue samples were collected with previous patient consent and in strict observance of legal and institutional ethical regulations. The protocol was approved by the French biomedical agency (Agence de la Biomédecine, approval number: PFS17-003).

### Human iPSC culture

The feeder-independent iPS cell line used for this study was a gift from Silvia Cappello (Max-Plank Institute of Psychiatry - Munich). Cells were reprogrammed from NuFF3-RQ human newborn foreskin feeder fibroblasts (GSC-3404, GlobalStel) (Kyrousi et al., 2021). iPS cells were cultivated as colonies on vitronectin-coated B3 dishes, using mTser medium (STEMCELL Technologies). Colonies were cleaned daily under a binocular stereo microscope (Lynx EVO, Vision engineering), by manually removing differentiated cells with a needle.

### Cerebral organoids culture

Cerebral organoids were derived from human iPS cells, following a previously published protocol (Qian et al., 2016). Day 0 to day 4: iPS colonies of 1-2 mm of diameter were detached with pre-warmed collagenase (1mg/mL) for 45 min at 37°C. After addition of 1 mL of mTser, floating colonies were transferred with a cut tip into a 15 ml tube for two series of gentle washing with medium 1 (DMEM-F12 without phenol red, 20% KOSR, 1X GlutaMAX, 1X MEM-NEAA, 1X 2-Mercaptoethanol, Pen/Strep, 2µM Dorsomorphin, 2 µM A-83). Colonies were subsequently distributed in an ultra-low attachment 6-well plate with 3 mL of medium 1 and cultivated at 37°C, 5% CO2. Day 5-6: Half of medium 1 was replaced daily with medium 2 (DMEM-F12 without phenol red, 1X N2 supplement, 1X GlutaMAX, 1X MEM-NEAA, Pen/Strep, 1 µM CHIR-99021, 1 µM SB-421542). Day 7-14: At day 7, EBs were embedded in Matrigel diluted in medium 2 at a ratio of 2:1. Matrigel-EB mixture was then spread in an ultra-low attachment dish and incubated at 37°C for 30 min to solidify (10-20 EBs per well). Finally, medium 2 was gently added to the well, without disturbing the Matrigel patch. At day 14, Matrigel was mechanically broken by pipetting with a 5 mL pipet and transferred into a 15 mL tube for gentle washing. Organoids were suspended in medium 3 (DMEM-F12 without phenol red, 1X N2 supplement, 1X b27 supplement, 1X GlutaMAX, 1X MEM-NEAA, 1X 2-Mercaptoethanol, Pen/Strep, 2.5 µg/mL Insulin) and grown in ultra-low attachment 6-well plates under agitation at 100 rpm (Digital Orbital Shaker DOS-10M from ELMI). Day 35 to 84: Starting from day 35, medium 3 was supplemented with diluted Matrigel (1:100) (Giandomenico et al., 2019).

### Infection of human fetal cortex and cerebral organoids

Fresh tissue from human fetal cortex was obtained from autopsies performed at the Robert Debré Hospital, and Necker enfants malades Hospital (Paris). A piece of pre-frontal cortex was collected from one hemisphere, and transported on ice from the hospital to the lab. The tissue was divided into smaller pieces and embedded 4% low-gelling agarose (Sigma) dissolved in artificial cerebrospinal fluid (ACSF). Cerebral organoids (week 8-12) were embedded in 3% low-gelling agarose. Gel blocks from both tissues were then sliced with a Leica VT1200S vibratome (300 µm-thick slices) in ice-cold ACSF. Slices were infected with a GFP coding retrovirus, diluted in DMEM-F12. After 2h of incubation, slices were washed three times with DMEM-F12 and grown on Millicell culture inserts (Merck) in cortical culture medium (DMEM-F12 containing B27, N2, 10 ng/ml FGF, 10 ng/ml EGF, 5% fetal bovine serum and 5% horse serum) for up to 5 days for human fetal brain and 48h for cerebral organoids. Medium was changed every day.

### Live imaging in cerebral organoids and human fetal cortex slices

To follow bRG cell divisions for approximatively 48h, we used the following approach. 48h after infection (3-5 days for human fetal brain), slices were placed under the microscope by transferring the culture inserts in a 35 mm FluoroDish (WPI) with 1 mL of culture medium (DMEM-F12 containing B27, N2, 10 ng/ml FGF, 10 ng/ml EGF, 5% fetal bovine serum and 5% horse serum). Live imaging was performed on a spinning disk wide microscope equipped with a Yokogawa CSU-W1 scanner unit to increase the field of view and improve the resolution deep in the sample. The microscope was equipped with a high working distance (WD 6.9-8.2 mm) 20X Plan Fluor ELWD NA 0.45 dry (Nikon), and a Prime95B SCMOS camera. Z-stacks of 80-100 µm range were taken with an interval of 4-5 µm, and maximum projections were performed. Videos were mounted in Metamorph. Image treatments (maximum projections, subtract background, Median filter, stackreg and rotation) were carried out on Fiji. Figures were assembled with Affinity Designer.

### Immunostaining of brain slices

Human fetal brain and cerebral organoid slices in culture were fixed in 4% PFA for 2 hours. Slices were boiled in sodium citrate buffer (10 mM, pH 6) for 20 minutes and cooled down at room temperature (antigen retrieval). Slices were then blocked in PBS-Triton 100X 0.3%-donkey serum 2% at room temperature for 2 hours, incubated with primary antibody overnight at 4°C in blocking solution, washed in PBS-Tween 0.05%, and incubated with secondary antibody overnight at 4°C in blocking solution before final wash and mounting in Aquapolymount. Mosaics (tilescans) of fixed tissue were acquired with a CFI Apo LWD Lambda S 40X objective (WI NA 1.15 WD 0.61-0.59, Nikon).

### Live and fixed correlative microscopy analysis

The correlative microscopy method enables to automatically pair and align live and fixed samples, for cell-cell matching. The macro, based on ImageJ (Schindelin et al., 2012) and Matlab, enables automated contouring of the slices, matching of the live and fixed samples based on their area and shape, and alignment of the samples (rotation and flip if needed). This leads to the precise positioning of the live imaged cells on the immunostained images. This method is described in detail in the **Annex 1**.

### Retrovirus production

To improve transfection efficiency, we used the HEK-Phoenix-GP cell line that stably expresses the packaging enzymes GAL and POL. Cells were plated in 3xT300 (dilution at 1:20) and grown 3 days to reach 70% of confluence in DMEM-GlutaMax medium, 10% FBS (50 mL/flask). At day 3, cells were transfected with envelope VSVG plasmid and transfer plasmid (CAG-GFP or MSCV-IRES-GFP) using Lipofectamine 2000. The two plasmids were mixed into 5.4 mL of OptiMEM medium (18 µg E-plasmid / 49.5 µg t-plasmid). 337.5 µL of Lipofectamine 2000 was diluted in 5.4 mL of OptiMEM medium and incubated 5 min at room temperature. The DNA preparation was thoroughly mixed into the Lipofectamine preparation and incubated 30 min at room temperature. In the meantime, medium was changed by 30 mL of DMEM-Glutamax (without FBS) per T300 flask. 3.6 mL of the DNA-Lipofectamine mixture was then added to each T300 flask and incubated 5h in a 37°C incubator. After this period, flasks were carefully transferred into an L3 lab and medium was changed for 30 mL of fresh DMEM-GlutaMAX, 10% FBS. At day 5, medium was harvested into 50 mL tubes and replaced by 30 mL of fresh medium (samples were stored at 4°C). At day 6, medium was harvested, pooled with Day 5 samples and spun-down to pellet cell debris (1300 rpm, 5 min at4°C). Supernatant was then filtered using 0.22 µm filter unit and divided into 6 Ultra-Clear tubs (Beckman Coulter – Ref.344058). Tubes were ultra-centrifuged at 31000 G for 1h30 at 4°C. Supernatant was removed, retroviruses were collected with multiple PBS washings and transferred into a single new Ultra-Clear tub. Final ultra-centrifugation was performed (31000 G for 1h30 at 4°C), supernatant was carefully removed and the thin pellet of retroviruses was suspended into 750 µL of DMEM-F12 medium, aliquoted (50 to 100 µL aliquots) and stored at -80°C. Titer of the preparation was tested by infecting regular HEK cells at different dilution and the percentage of GFP+ cells was measured by FACS.

### Expression constructs and antibodies

The following plasmids were used in this study: CAG-GFP (a gift from Victor Borrell); MSCV-IRES-GFP (Tannishtha Reya, Addgene 20672); VSVG (a gift from Philippe Benaroch). Antibodies used in this study were mouse anti-SOX2 (Abcam Ab79351, 1/500), sheep anti-EOMES (R&D Sytems AF6166, 1/500), rabbit anti-NEUN (Abcam Ab177487, 1/500), chicken anti-GFP (Abcam Ab13970, 1/500), mouse anti-pVimentin (Abcam Ab22651, 1/1000), rat anti-HES1 (MBL D134-3, 1/500).

## Supporting information

Annexe 1

## Acknowledgements

We acknowledge Institut Curie, member of the French National Research Infrastructure France-BioImaging (ANR10-INBS-04) and the Nikon BioImaging Center (Institut Curie, France). We thank Fiona Francis (IFRM) and Xavier Morin for helpful discussions and critical reading of the manuscript. L.C. was funded by the French ministry of research (MESRI). A-S.M. was funded by the Labex Cell(n)Scale and CNRS. A.D.B. is an INSERM researcher. This work was supported by the CNRS, I. Curie, the ANR (ANR-20-CE16-0004-01) and the Ville de Paris “Emergences” program.

## Author contributions

L.C. performed experiments, analyzed data and wrote the manuscript. A-S.M. coded the LiveFixedCorrelative macro, C.B.A. analyzed data, S.F., A.D.C. and M.L. generated organoids, B.S, T.A-B. and F.G. provided fetal tissue, V.F. assisted with imaging and A.D.B. designed the project and wrote the manuscript.

---

**Figure S1.**
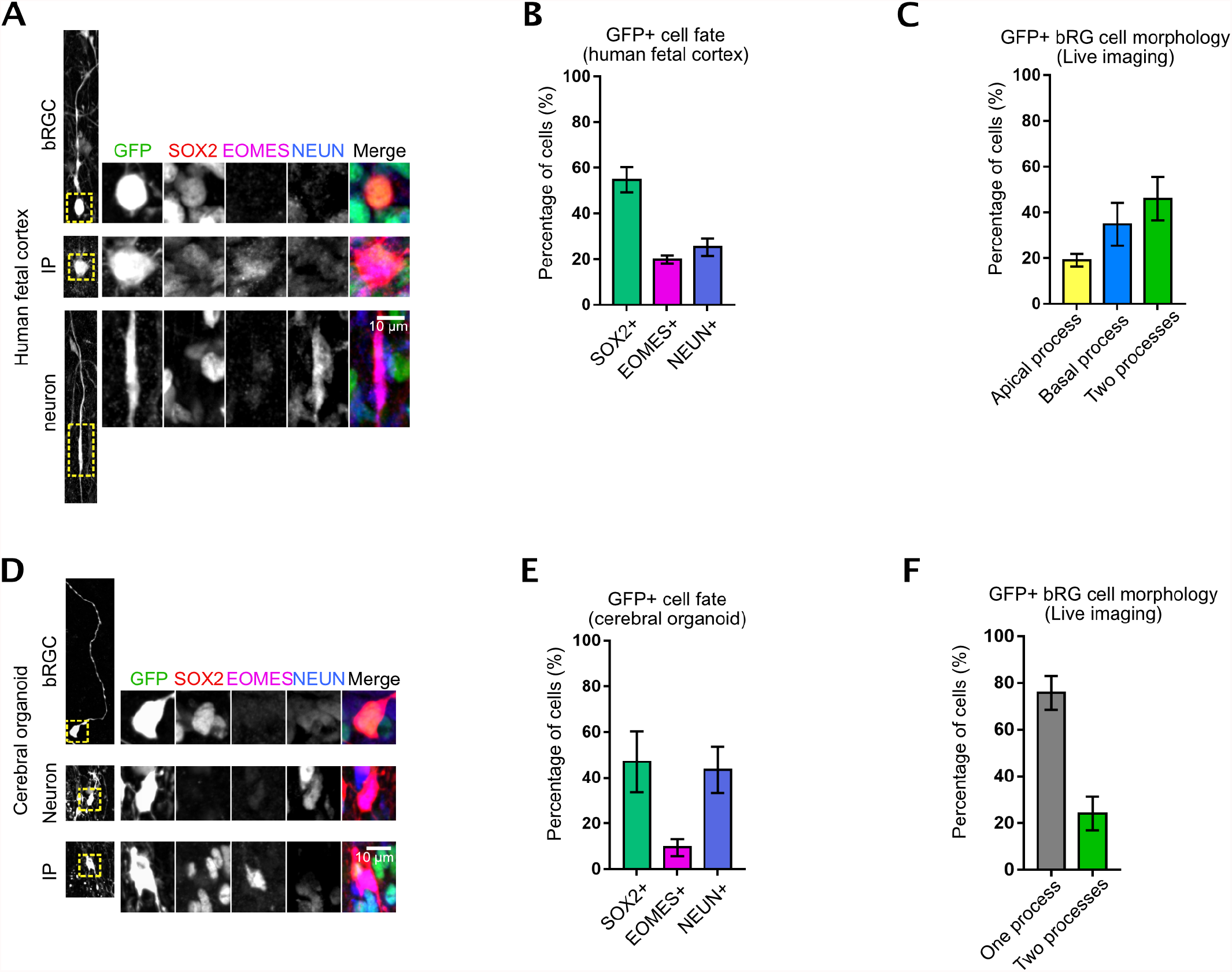
Fate and shape of GFP+ cells in cerebral organoids and fetal tissue. **A**. Immunostaining for SOX2, EOMES and NEUN in GFP-infected human fetal cortex at GW 17. **B**. Fate of GFP+ cells in human fetal cortex at GW 14-18. **C**. Morphology of GFP+ bRG cells in live imaged human fetal samples at GW 14-18. **D**. Immunostaining for SOX2, EOMES and NEUN in GFP-infected cerebral organoids at week 8. **E**. Fate of GFP+ cells in cerebral organoids at week 8-10. **F**. Morphology of GFP+ bRG cells in live imaged cerebral organoids at week 8-10.

**Figure S2.**
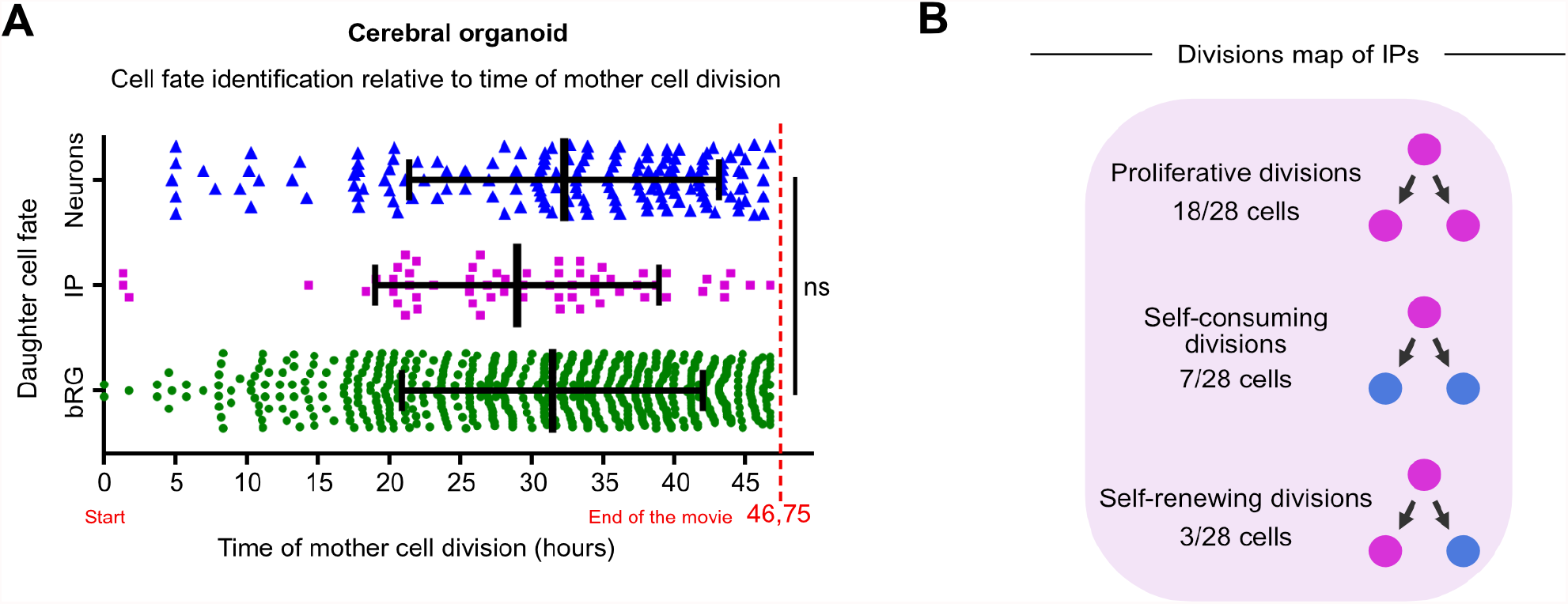
Cell fate identification timing and IP division modes in cerebral organoids. **A**. Detection of bRG, IP or neuronal cell fate relative to the time of division of the bRG mother cell in cerebral organoids at week 8-10. **B**. Summary of division patterns identified in IPs within week 8-10 cerebral organoids (N=28 IPs).

**Figure S3.**
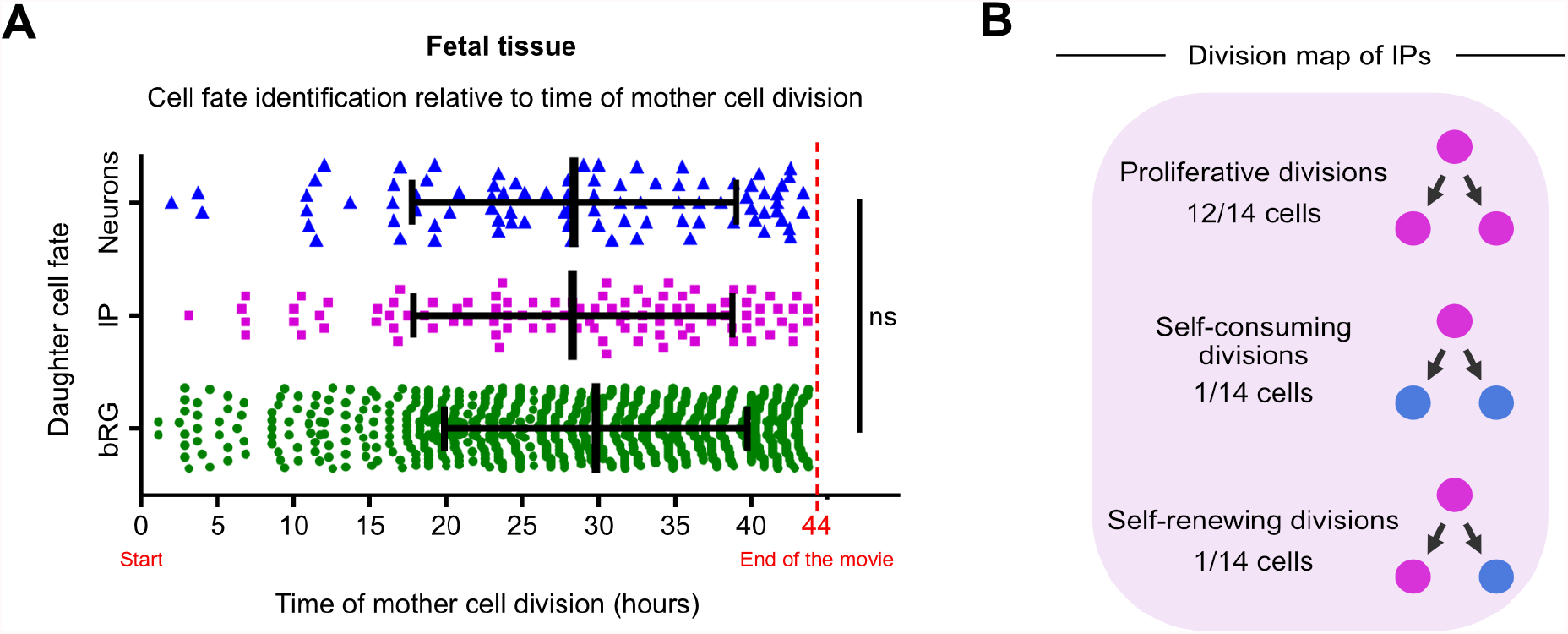
Cell fate identification timing and IP division modes in fetal tissue. **A**. Detection of bRG, IP or neuronal cell fate relative to the time of division of the bRG mother cell in human fetal samples at GW 14-18. **B**. Summary of division patterns identified in IPs within human fetal samples at GW 14-18 (N=14 IPs).

**Figure S4.**
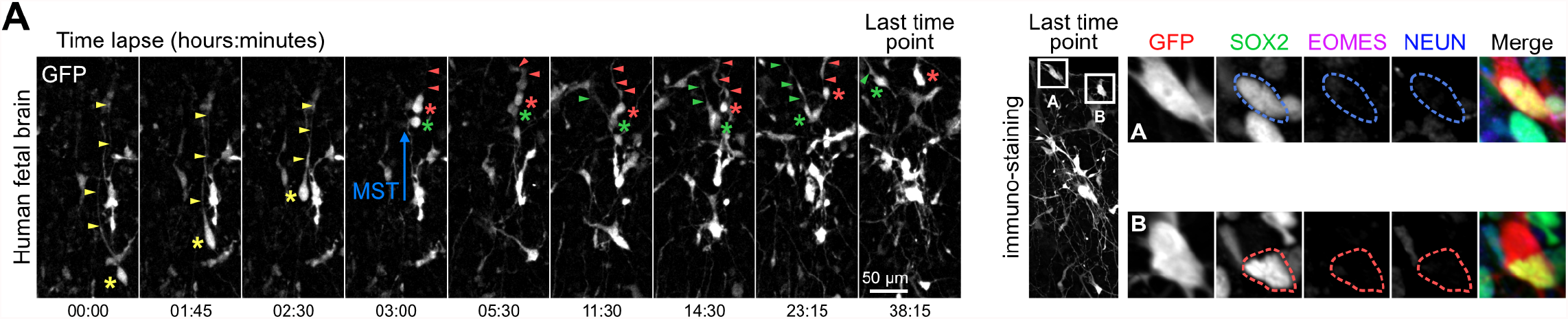
SOX2+ bRG daughter cells regrow a basal process if at birth. **A**. Live/fixed correlative analysis of a dividing bRG cell generating two bRG daughters. Asterix indicates cell soma and arrowhead indicates basal process. Mother cell (yellow) divides into a process-inheriting cell (red) and a cell that regrows a basal process (green).

