## Supplementary material for "A cell fate decision map reveals abundant direct neurogenesis in the human developing neocortex": Annexe 1

### Documentation for the LiveFixedCorrelative code

#### Table of contents

1. Context of the macro ... 3
1.1. Image acquisition ... 3
1.2. Separating biological conditions ... 4
1.3. Input of the macro ... 4
2. STG file analysis and inputs for the macro ... 5
3. Chosen file architecture ... 6
Errors linked to a default of folder architecture: ... 6
4. First use of the macro ... 8
Installation of the executable MATLAB ... 8
5. Global presentation of the steps: ... 9
5.1. Acquisition and analysis parameters, folder choice ... 9
5.2. Condition choice ... 10
5.3. Creation of the information file for MATLAB ... 10
5.4. Reconstruction of the multi-stage acquisition ... 10
5.5. (Manual) segmentation of the 1st modality reconstructed image ... 11
5.6. (Automatic) segmentation of the 2nd modality images ... 12
5.7. Representation of the movie acquisitions on the reconstruction ... 13
5.8. Pairing of the matching structures ... 13
5.9. Alignment of the paired structures ... 14
5.10. Application of the transformation on images ... 14
6. Detailed presentation of the three interfaces ... 15
6.1. Presentation of the ImageJ interface ... 15
**6.1.1. Information about the live acquisition & 2/ Information about fix images ... 15**
**6.1.2. Segmentation on the fix mosaic images ... 15**
**6.1.3. Saving results ... 15**
6.2. Presentation of the first MATLAB interface: pairing of the contours ... 17
**6.2.1. Display and visibility of contours ... 18**
**6.2.2. Association of the contours ... 18**
**6.2.3. Duplicate contours ... 18**
**6.2.4. Show images ... 18**
**6.2.5. Validate ... 19**

### 1. Context of the macro

The live and fixed correlative analysis enables to follow dynamic events such as mitosis or cell migration in order to analyze specific features of these cells or their progeny. The difficulty of such analysis is to be able to find the imaged cells once the tissue is fixed, especially if multiple slices of tissue (from different conditions) were imaged at the same time. Following fixation, immunostaining and mounting, slices of tissues do not remain aligned with their orientation in the movie. It is therefore very difficult to find corresponding areas within these movies and fixed acquisitions. We therefore developed a contour-based approach that enables to pair live and fixed images, align them together and identify each live imaging position in the images acquired after fixation and staining.

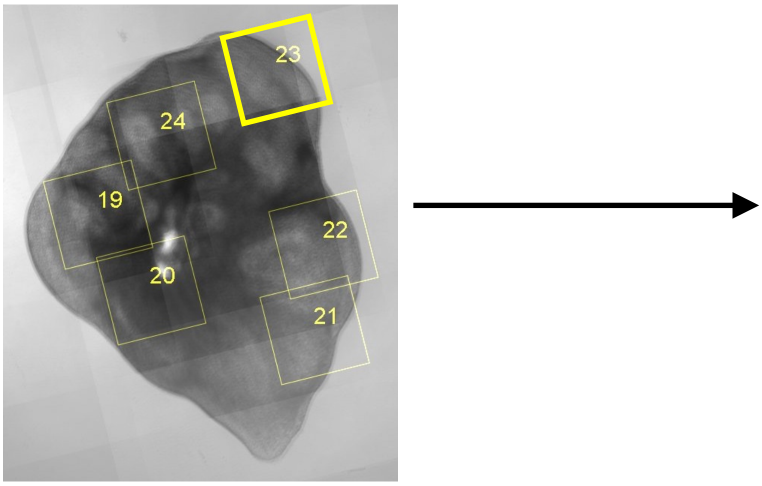

4X Transmitted light (Live)

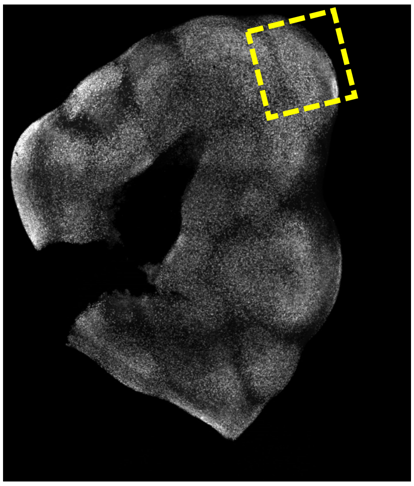

10X mosaic (Immunostaining)

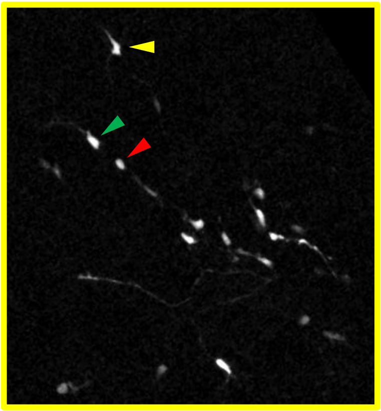

Last time point of the movie

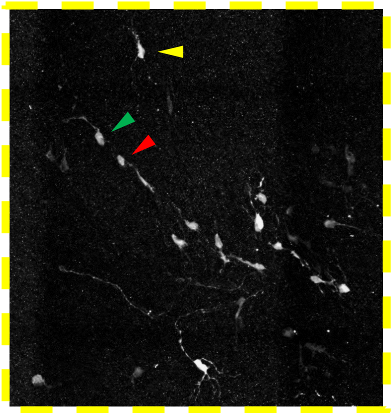

40X mosaic (Immunostaining)

The main task of the code is to find the transformation between the contours of the tissues imaged in both Live and Fixed modalities. This process is very complex and therefore requires a user-friendly interface that offers the possibility, if necessary, to perform manual adjustment at each step of the workflow.

#### 1.1. Image acquisition

In our case, the images were acquired using different modalities:

- First modality: a set of images acquired in transmitted light microscopy, as a multi-stage acquisition
    - ➔ A reconstruction is required to see the whole tissue

  *Remark: the stage positions correspond to stage positions where movies were acquired; some non-acquired positions can be added to obtain a more precise reconstruction of the whole structure. In our case, these images are acquired at the end of the live acquisition, with an objective of smaller magnification*.

- Second modality: images after fixation acquired in fluorescence microscopy, using a mosaic mode. Each image contains one slice of tissue. In our case, mosaics were reconstructed with Imaris Stitcher software (Bitplane) v.9.5.0, and maximum z-projection is used.

##### 1.2. Separating biological conditions

Because the slices corresponding to different biological conditions were placed in different dishes (up to four), we decided to separate the treatment of these conditions. Therefore, a reconstruction of the transmitted light images is performed, using the stage of each position within the same condition (or dish).
*Remark: This separation is necessary to create images of reasonable size, easy to handle afterwards.*

Several slices of tissues are live-imaged and the condition corresponding to each image is specified as follows:

- For the first modality: within the name chosen for each stage position (see here)
- For the second modality: in the folder name containing the second modality images, named after the condition (see the architecture description here)

##### 1.3. Input of the macro

- Images in the first modality (referring to “Live”, “transmitted light images”, or “movie” in this documentation) result in a multi-stage acquisition, with a ND file enabling to read them all together.
- Images in the second modality (referring to “Fixed images”, “fluorescence” or “mosaic” in this documentation) result in one-plane images (the format is not important, but TIF is recommended)
- A “STG” file, generated by Metamorph, that gives the stage positions and associates a name at each image of the first modality

#### 2. STG file analysis and inputs for the macro

The STG file is generated by Metamorph when a multi-stage acquisition is performed; it concerns the first modality images. Two meaningful information are read from this file:

- The position of the stage for each image, (X,Y) coordinates in µm,
- The name associated to each stage position.

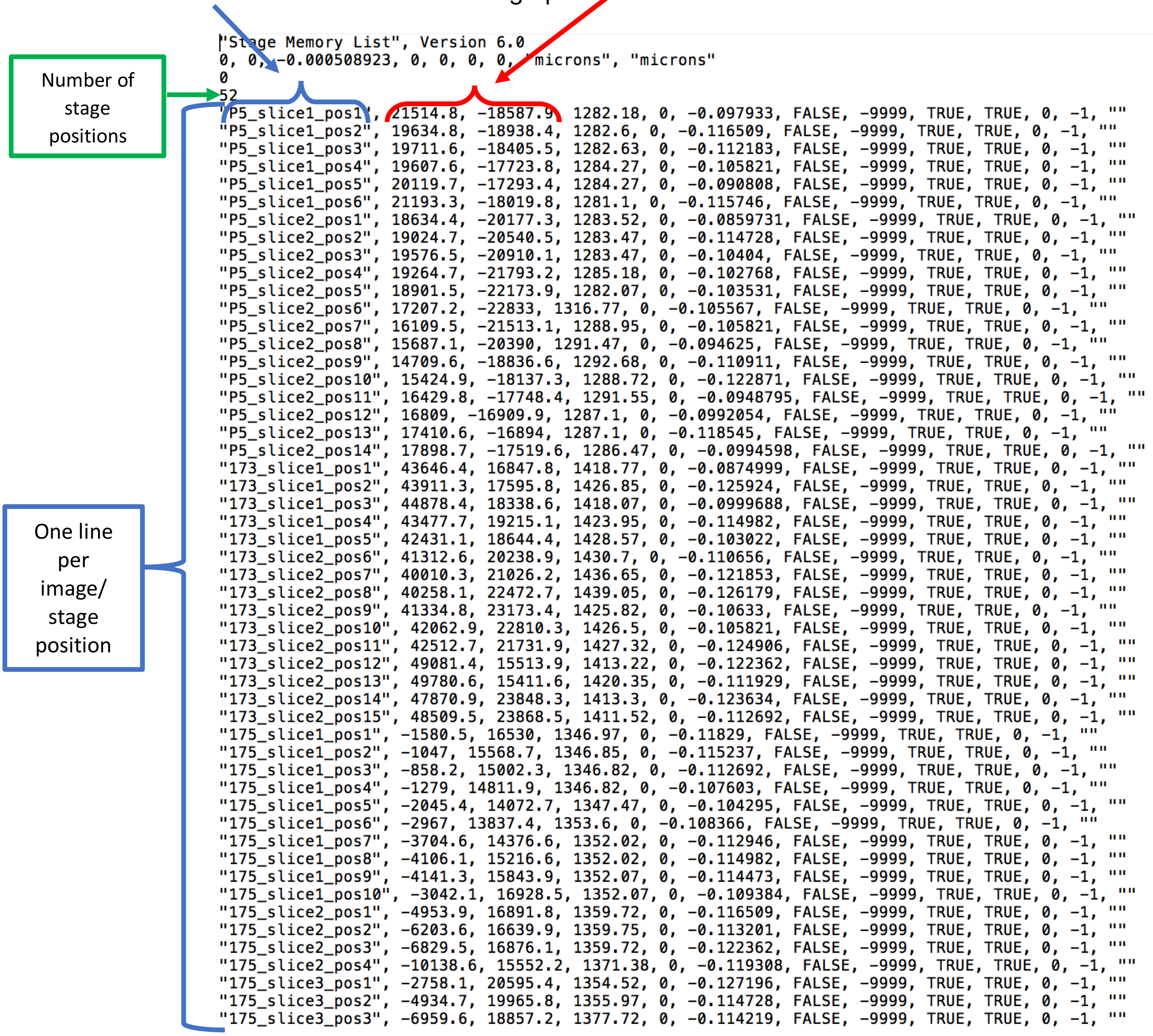

```
"Stage Memory List", Version 6.0
0, 0, -0.000508923, 0, 0, 0, 0, "microns", "microns"
0
52
"P5_slice1_pos1", 21514.8, -18587.9, 1282.18, 0, -0.097933, FALSE, -9999, TRUE, TRUE, 0, -1, ""
"P5_slice1_pos2", 19634.8, -18938.4, 1282.6, 0, -0.116509, FALSE, -9999, TRUE, TRUE, 0, -1, ""
"P5_slice1_pos3", 19711.6, -18405.5, 1282.63, 0, -0.112183, FALSE, -9999, TRUE, TRUE, 0, -1, ""
"P5_slice1_pos4", 19607.6, -17723.8, 1284.27, 0, -0.105821, FALSE, -9999, TRUE, TRUE, 0, -1, ""
"P5_slice1_pos5", 20119.7, -17293.4, 1284.27, 0, -0.090808, FALSE, -9999, TRUE, TRUE, 0, -1, ""
"P5_slice1_pos6", 21193.3, -18019.8, 1281.1, 0, -0.115746, FALSE, -9999, TRUE, TRUE, 0, -1, ""
"P5_slice2_pos1", 18634.4, -20177.3, 1283.52, 0, -0.0859731, FALSE, -9999, TRUE, TRUE, 0, -1, ""
"P5_slice2_pos2", 19024.7, -20540.5, 1283.47, 0, -0.114728, FALSE, -9999, TRUE, TRUE, 0, -1, ""
"P5_slice2_pos3", 19576.5, -20910.1, 1283.47, 0, -0.10404, FALSE, -9999, TRUE, TRUE, 0, -1, ""
"P5_slice2_pos4", 19264.7, -21793.2, 1285.18, 0, -0.102768, FALSE, -9999, TRUE, TRUE, 0, -1, ""
"P5_slice2_pos5", 18901.5, -22173.9, 1282.07, 0, -0.103531, FALSE, -9999, TRUE, TRUE, 0, -1, ""
"P5_slice2_pos6", 17207.2, -22833, 1316.77, 0, -0.105567, FALSE, -9999, TRUE, TRUE, 0, -1, ""
"P5_slice2_pos7", 16109.5, -21513.1, 1288.95, 0, -0.105821, FALSE, -9999, TRUE, TRUE, 0, -1, ""
"P5_slice2_pos8", 15687.1, -20390, 1291.47, 0, -0.094625, FALSE, -9999, TRUE, TRUE, 0, -1, ""
"P5_slice2_pos9", 14709.6, -18836.6, 1292.68, 0, -0.110911, FALSE, -9999, TRUE, TRUE, 0, -1, ""
"P5_slice2_pos10", 15424.9, -18137.3, 1288.72, 0, -0.122871, FALSE, -9999, TRUE, TRUE, 0, -1, ""
"P5_slice2_pos11", 16429.8, -17748.4, 1291.55, 0, -0.0948795, FALSE, -9999, TRUE, TRUE, 0, -1, ""
"P5_slice2_pos12", 16809, -16909.9, 1287.1, 0, -0.0992054, FALSE, -9999, TRUE, TRUE, 0, -1, ""
"P5_slice2_pos13", 17410.6, -16894, 1287.1, 0, -0.118545, FALSE, -9999, TRUE, TRUE, 0, -1, ""
"P5_slice2_pos14", 17898.7, -17519.6, 1286.47, 0, -0.0994598, FALSE, -9999, TRUE, TRUE, 0, -1, ""
"173_slice1_pos1", 43646.4, 16847.8, 1418.77, 0, -0.0874999, FALSE, -9999, TRUE, TRUE, 0, -1, ""
"173_slice1_pos2", 43911.3, 17595.8, 1426.85, 0, -0.125924, FALSE, -9999, TRUE, TRUE, 0, -1, ""
"173_slice1_pos3", 44878.4, 18338.6, 1418.07, 0, -0.0999688, FALSE, -9999, TRUE, TRUE, 0, -1, ""
"173_slice1_pos4", 43477.7, 19215.1, 1423.95, 0, -0.114982, FALSE, -9999, TRUE, TRUE, 0, -1, ""
"173_slice1_pos5", 42431.1, 18644.4, 1428.57, 0, -0.103022, FALSE, -9999, TRUE, TRUE, 0, -1, ""
"173_slice2_pos6", 41312.6, 20238.9, 1430.7, 0, -0.110656, FALSE, -9999, TRUE, TRUE, 0, -1, ""
"173_slice2_pos7", 40010.3, 21026.2, 1436.65, 0, -0.121853, FALSE, -9999, TRUE, TRUE, 0, -1, ""
"173_slice2_pos8", 40258.1, 22472.7, 1439.05, 0, -0.126179, FALSE, -9999, TRUE, TRUE, 0, -1, ""
"173_slice2_pos9", 41334.8, 23173.4, 1425.82, 0, -0.10633, FALSE, -9999, TRUE, TRUE, 0, -1, ""
"173_slice2_pos10", 42062.9, 22810.3, 1426.5, 0, -0.105821, FALSE, -9999, TRUE, TRUE, 0, -1, ""
"173_slice2_pos11", 42512.7, 21731.9, 1427.32, 0, -0.124906, FALSE, -9999, TRUE, TRUE, 0, -1, ""
"173_slice2_pos12", 49081.4, 15513.9, 1413.22, 0, -0.122362, FALSE, -9999, TRUE, TRUE, 0, -1, ""
"173_slice2_pos13", 49780.6, 15411.6, 1420.35, 0, -0.111929, FALSE, -9999, TRUE, TRUE, 0, -1, ""
"173_slice2_pos14", 47870.9, 23848.3, 1413.3, 0, -0.123634, FALSE, -9999, TRUE, TRUE, 0, -1, ""
"173_slice2_pos15", 48509.5, 23868.5, 1411.52, 0, -0.112692, FALSE, -9999, TRUE, TRUE, 0, -1, ""
"175_slice1_pos1", -1580.5, 16530, 1346.97, 0, -0.11829, FALSE, -9999, TRUE, TRUE, 0, -1, ""
"175_slice1_pos2", -1047, 15568.7, 1346.85, 0, -0.115237, FALSE, -9999, TRUE, TRUE, 0, -1, ""
"175_slice1_pos3", -858.2, 15002.3, 1346.82, 0, -0.112692, FALSE, -9999, TRUE, TRUE, 0, -1, ""
"175_slice1_pos4", -1279, 14811.9, 1346.82, 0, -0.107603, FALSE, -9999, TRUE, TRUE, 0, -1, ""
"175_slice1_pos5", -2045.4, 14072.7, 1347.47, 0, -0.104295, FALSE, -9999, TRUE, TRUE, 0, -1, ""
"175_slice1_pos6", -2967, 13837.4, 1353.6, 0, -0.108366, FALSE, -9999, TRUE, TRUE, 0, -1, ""
"175_slice1_pos7", -3704.6, 14376.6, 1352.02, 0, -0.112946, FALSE, -9999, TRUE, TRUE, 0, -1, ""
"175_slice1_pos8", -4106.1, 15216.6, 1352.02, 0, -0.114982, FALSE, -9999, TRUE, TRUE, 0, -1, ""
"175_slice1_pos9", -4141.3, 15843.9, 1352.07, 0, -0.114473, FALSE, -9999, TRUE, TRUE, 0, -1, ""
"175_slice1_pos10", -3042.1, 16928.5, 1352.07, 0, -0.109384, FALSE, -9999, TRUE, TRUE, 0, -1, ""
"175_slice2_pos1", -4953.9, 16891.8, 1359.72, 0, -0.116509, FALSE, -9999, TRUE, TRUE, 0, -1, ""
"175_slice2_pos2", -6203.6, 16639.9, 1359.75, 0, -0.113201, FALSE, -9999, TRUE, TRUE, 0, -1, ""
"175_slice2_pos3", -6829.5, 16876.1, 1359.72, 0, -0.122362, FALSE, -9999, TRUE, TRUE, 0, -1, ""
"175_slice2_pos4", -10138.6, 15552.2, 1371.38, 0, -0.119308, FALSE, -9999, TRUE, TRUE, 0, -1, ""
"175_slice3_pos1", -2758.1, 20595.4, 1354.52, 0, -0.127196, FALSE, -9999, TRUE, TRUE, 0, -1, ""
"175_slice3_pos2", -4934.7, 19965.8, 1355.97, 0, -0.114728, FALSE, -9999, TRUE, TRUE, 0, -1, ""
"175_slice3_pos3", -6959.6, 18857.2, 1377.72, 0, -0.114219, FALSE, -9999, TRUE, TRUE, 0, -1, ""
```

These values are used to compute:

- The *translation of the stage* in **pixels**, according to the first position of the condition.

- The *condition name, associated to each stage of the first modality*. In our case, we decided to use the string at the beginning of the position name, before "_slice".

*Remark: The delimiter to separate condition is totally arbitrary and can be easily changed, see here. The images corresponding to the stage positions to reconstruct together must have a common name, easy to extract from the position names in the STG file.*

#### 3. Chosen file architecture

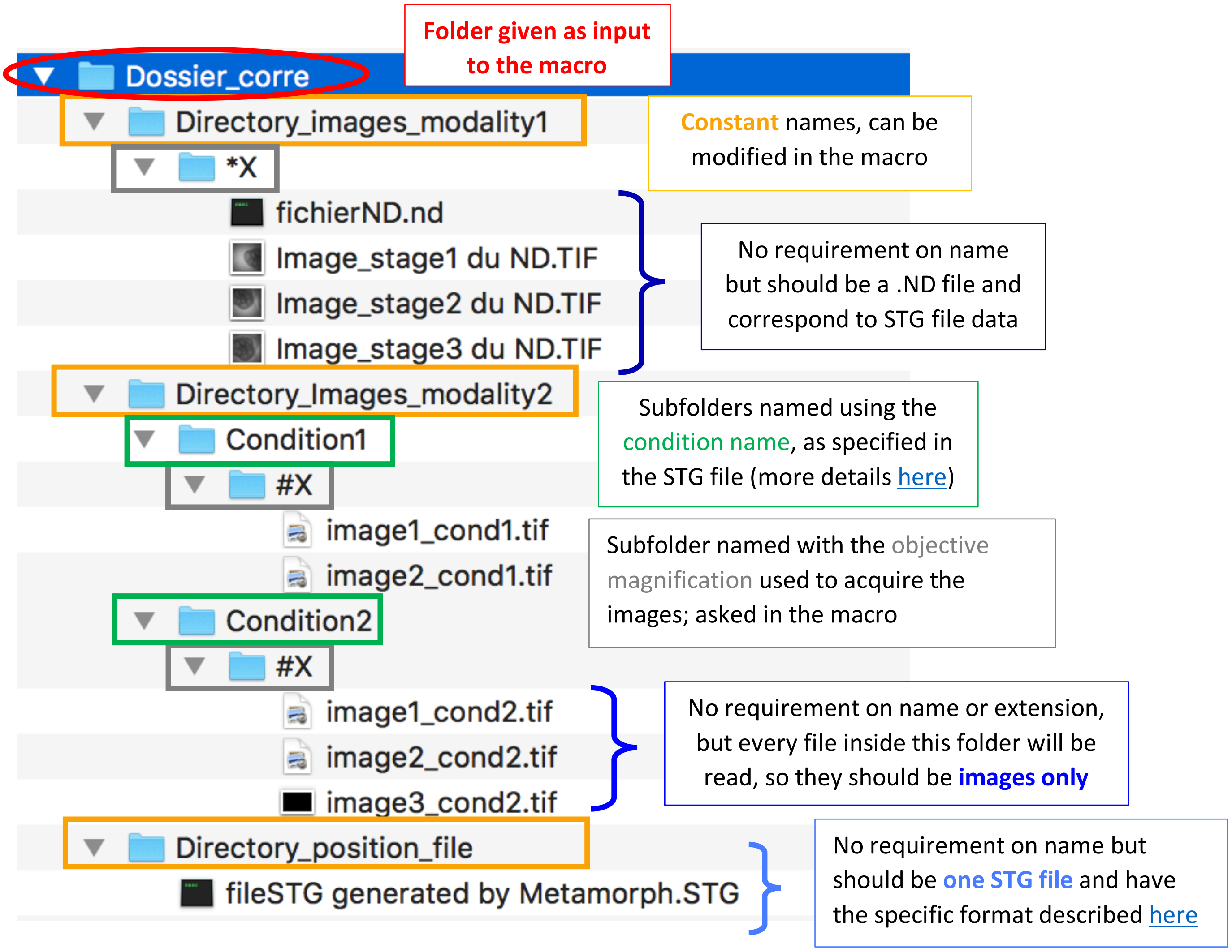

Do not hesitate to refer to the example data set for a better understanding of the architecture.

##### Errors linked to a default of folder architecture:

If the folder architecture is not respected: some error messages appears (and the macro stops):

- Make sure there is no space in the full name of the folder (calling the MATLAB exe with space in the name will not work because space is a delimiter to specify arguments to the executable of MATLAB).

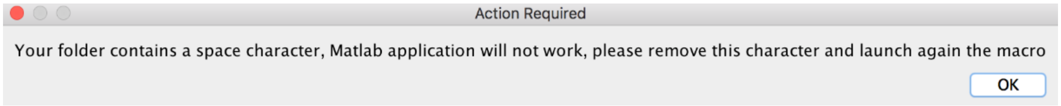

- Only images should be in the second modality sub-folders, otherwise an error "There are no image open" occurs and the macro stops. It is possible to create subfolder(s), the macro ignores them (with different name than "Results", this one is created during the macro and contains the contours).

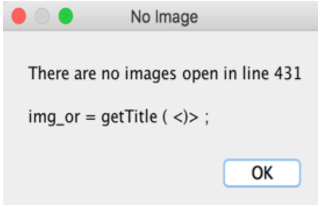

- Concerning the STG folder: either no folder named as *Directory_position_file* or no STG file inside:

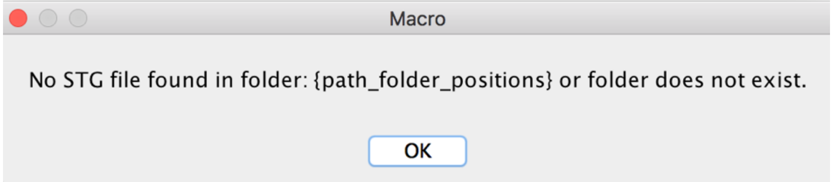

- Concerning the first modality: either no folder with the specified name *Directory_images_modality1* or no subfolder containing the data with the correct name (magnification-based name):

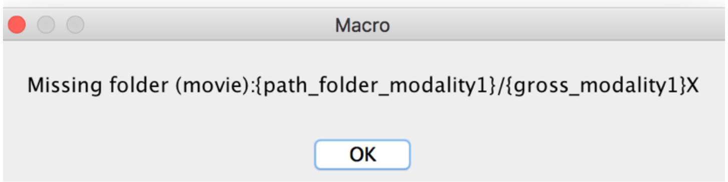

- Concerning the second modality: either no folder found with the specified name *Directory_images_modality2* or no sub-(sub-)folder containing the data with the correct name (folders with condition-based names, containing each one a sub-folder with magnification-based name):

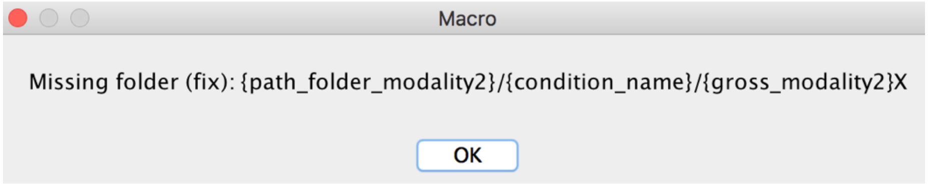

Some other errors can appear:

- No ND file corresponding to modality 1:

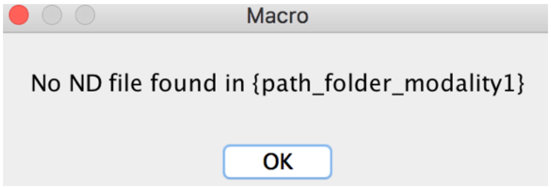

- More than one ND file corresponding to modality 1:

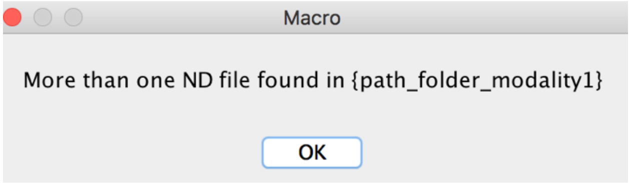

- Several STG files in the folder *Directory_position_file*:

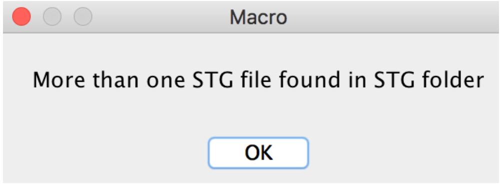

### 4. First use of the macro

#### Installation of the executable MATLAB

Follow the steps below:

- Run as an administrator the executable *correlative_MATLABinterface_exe*:

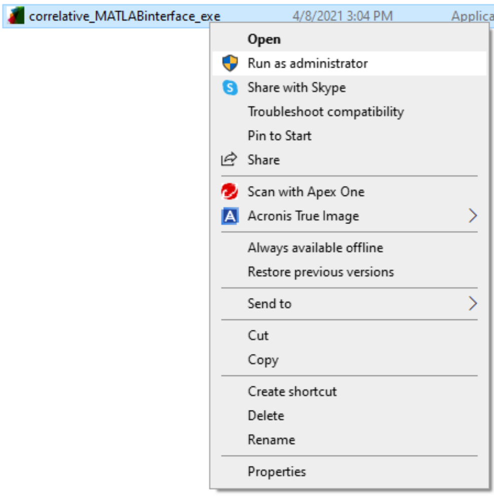

- The installation window will appear, follow the steps. The installation should be done in a folder named "CorrelativeExe" that you will create in the Plugin folder of Fiji:

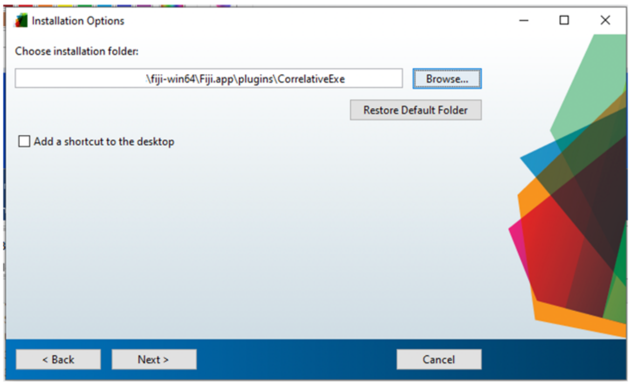

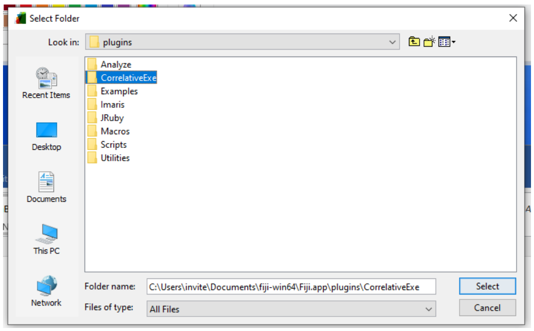

This program contains both the exe called by the macro but also the MATLAB Runtime (https://www.mathworks.com/products/compiler/matlab-runtime.html) necessary to make the exe run; you can install this part in the default proposed folder.

*Remark: you can also install the executable in another location and directly give the (absolute) path to the macro, line 8:*

```
7    // path of the matlab exe
8    exe_path = getDirectory("plugins")+"CorrelativeExe"+File.separator+"application"+File.separator+"correlation_contours_trans_fluo.exe";
```

To run the macro, either Drag and drop it and click on "Run" or put it in the "Plugin" folder of ImageJ and run it via Plugins **LiveFixedCorrelative**.

As explained before, the architecture of the folders/files is very important to make the macro work. At the beginning of the macro (lines 14-16), we offer the possibility to change the folder names; please refer to the example data set if you have a doubt:

```
// folder organization
directory_images_modality1  = "trans/";
directory_images_modality2 = "fix/"
directory_position_file = "STG/";
```

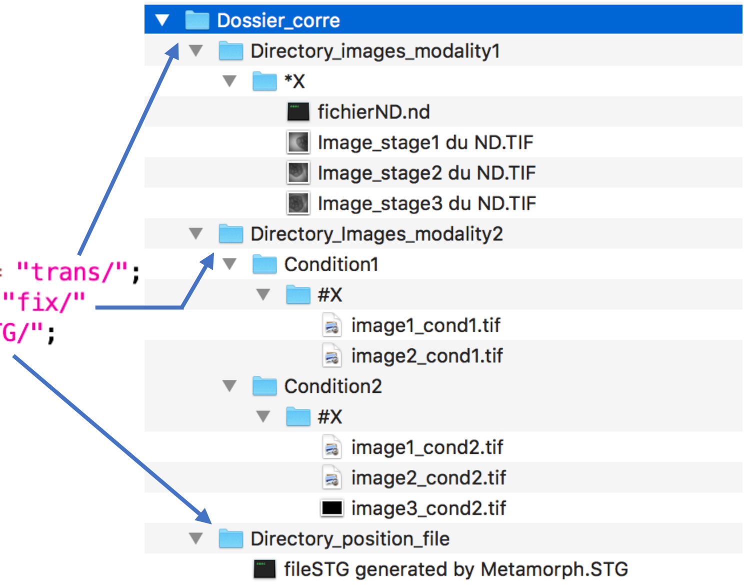

#### 5. Global presentation of the steps:

- Acquisition and parameter analysis, folder choice
- Condition choice
- Creation of the information file for MATLAB
- Reconstruction of the trans acquisition using different stage positions
- (Manual) segmentation of the trans reconstruction
- (Automatic) segmentation of the 2nd modality images
- Representation of the movie acquisitions on the reconstruction
- Pairing of the matching structures
- Alignment of the pairing structures
- Application of the transformation on images

##### 5.1. Acquisition and parameter analysis, folder choice

First the user should specify the acquisition parameters. This step is important because the parameters define the name of the subfolders where the inputs should be located, see the full description):

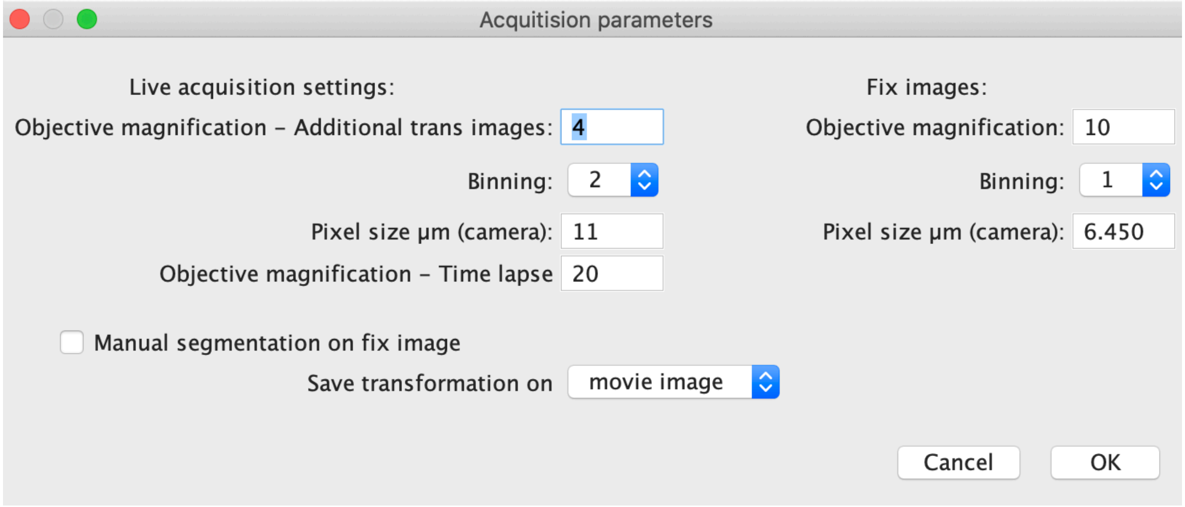

Then the user is asked to load a folder; this folder must respect the requirements defined here and will be called in the following *main directory* or *input directory*.

*Remark: In the given example, a folder named "4X" is created for the first modality and a folder "10X" is created in each sub-folder of conditions for the second modality.*

#### 5.2. Condition choice

The macro offers the possibility to treat only some of the biological conditions (conditions as they are defined in the STG). → Check only the ones you want to treat.

#### 5.3. Creation of the information file for MATLAB

This step is invisible for the user but important because it generates a "communication file" between the ImageJ macro and the MATLAB program. This file is called *recap_for_matlab.txt* and saved in a folder named *InfoForMatlab*, created by the macro. It provides the names of the folders containing images for each modality, all the parameters to determine the pixel size of each image and which conditions to treat:

| | Name | Value |
|---|---|---|
| 1 | Directory_Fixed_Images | {chosen_directory_path}/fix/ |
| 2 | Directory_Movie_Images | {chosen_directory_path}/trans/4X |
| 3 | Percentage_subsampling | 0.250 |
| 4 | Movie_objective | 4 |
| 5 | Movie_binning | 2 |
| 6 | Movie_pixel_size | 11.000 |
| 7 | Fix_objective | 10 |
| 8 | Fix_binning | 1 |
| 9 | Fix_pixel_size | 6.450 |
| 10 | Number of conditions | 1 |
| 11 | Condition1 | P5 |

#### 5.4. Reconstruction of the multi-stage acquisition

The images of the first modality are acquired as a multi-stage acquisition. To distinguish the whole structures, a bigger Z-stack image is created in which each slice corresponds to one image, translated according to the stage position given in the STG file. Then a standard-deviation Z-projection is performed on the stack to obtain the final reconstructed image.

*Remark: only images corresponding to a same condition are reconstructed together. The first image of the condition is centered in this big image.*

Basic example with four images:

- Trans separated images

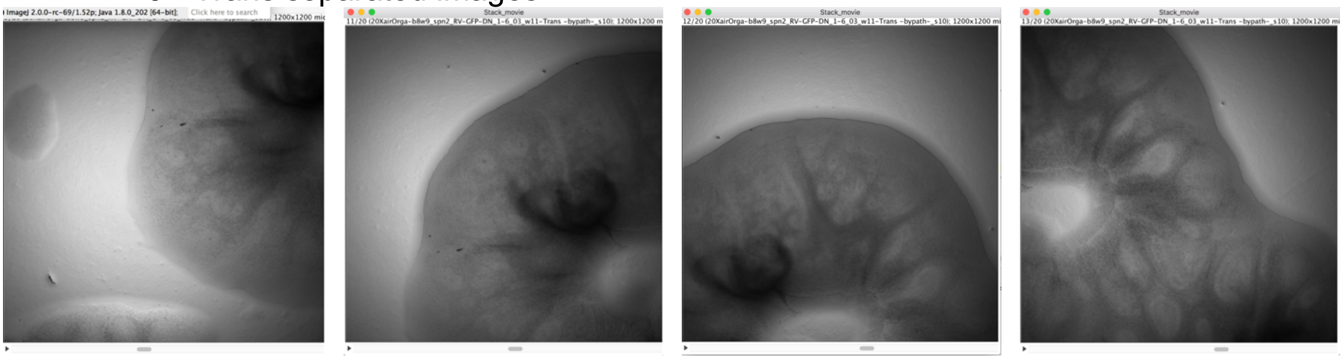

- Stage movement:

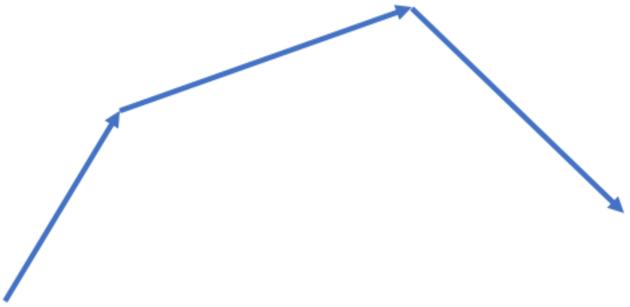

- Translated images in a bigger image (size computed to contain **<u>all</u>** images of the condition), (organized as a Stack in ImageJ)

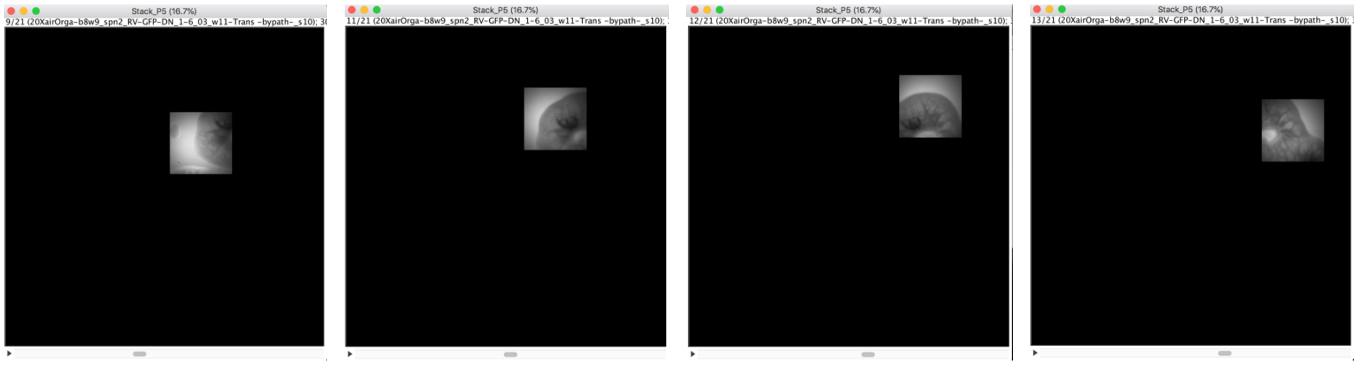

- Standard deviation of the stack: structures can be distinguished

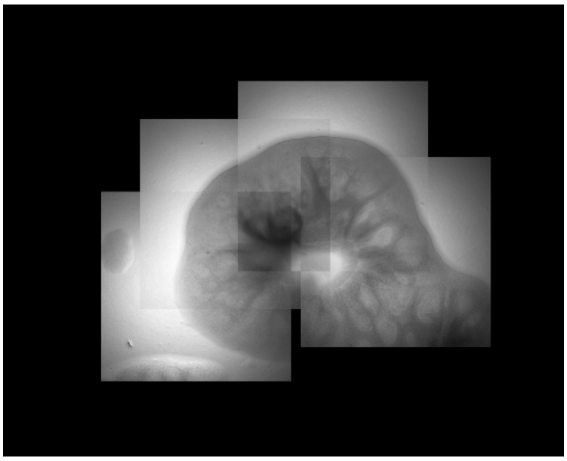

##### 5.5. (Manual) segmentation of the 1st modality reconstructed image

In our case, the first modality images were not easy to segment automatically (transmitted light images & multi-stage acquisition creating images with different dynamics). We opted for a manual segmentation with the tool "Polygon selection", where the user has to add each shape he wants to study in the ROI Manager:

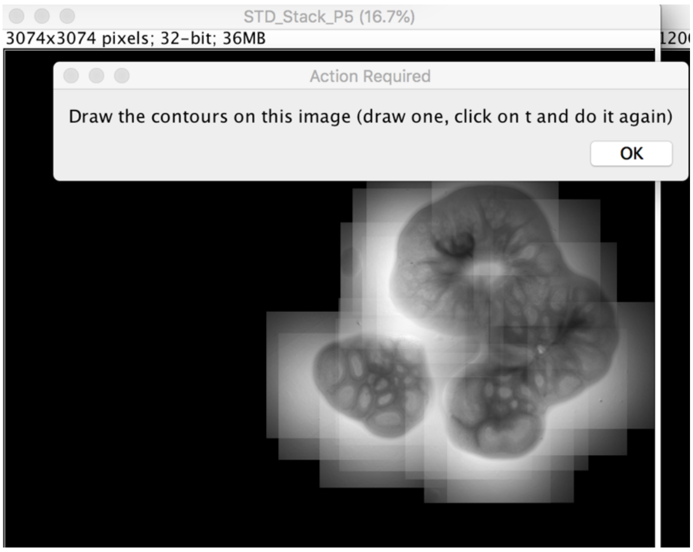

These drawn ROIs are saved in a folder "*ROI*" created by the macro within the main directory, under the name "*ConditionName*_RoiSet.zip". If the macro is relaunched on the same set, ROIs will be automatically loaded and displayed on the image, with the possibility to modify/delete them:

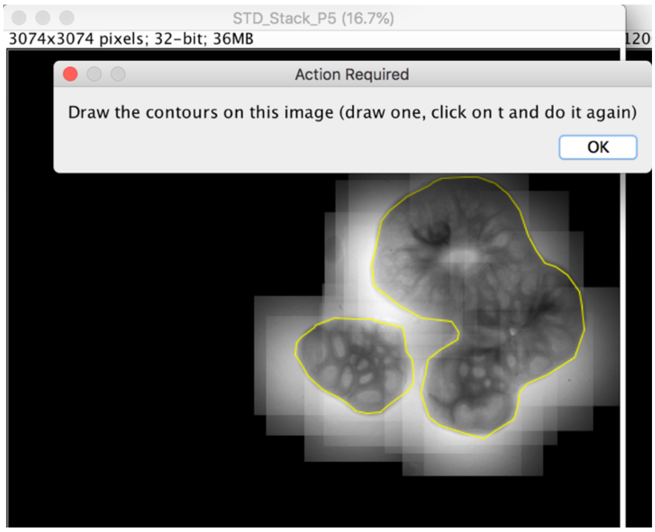

The contour of each tissue (one ROI representing one tissue) is saved as a set of coordinates in a sub-folder *Results*, created by the macro, within the folder containing the images. A crop on each shape is also saved for the MATLAB program.

#### 5.6. (Automatic) segmentation of the $2^{nd}$ modality images

This segmentation has been optimized based on our images, containing only one slice of tissue per image (here is how to change it). The steps are:

- Subsample the image (factor 4)
- Subtract Background (rolling ball of 100 pixels)
- Convert in 8 bits
- Threshold using Percentile method
- Remove outliers (radius 2)
- Perform 10 iterations of dilation
- Fill holes
- Perform 10 iterations of Erode
- Keep biggest ROI

For more flexibility, the macro offers a “Manual mode” where the user can draw the contours himself on the different images.

Contours of the tissue from each image is saved as a set of coordinates in a sub-folder created by the macro named *Results* located in the folder containing the images.

#### 5.7. Representation of the movie acquisitions on the reconstruction

In order to retrieve the cells imaged during the movie, the location of each recorded movie is required. Squares representing the movie fields of view are displayed on the reconstruction with the corresponding stage number:

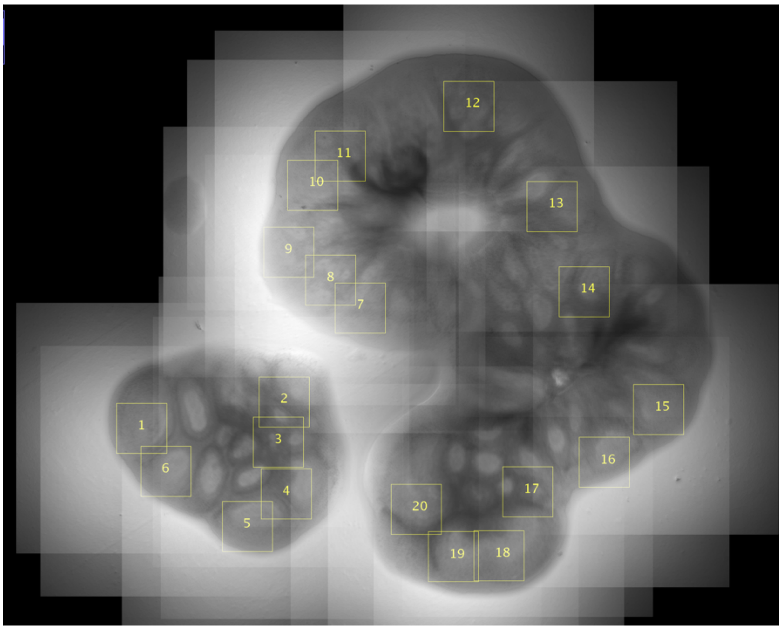

This image with squares is saved in the folder *Results* located in the main directory, under the name: “Composite-*ConditionName*.tif”.

*Remark: to locate these positions on this reconstruction with a great accuracy, you must check that the microscope is paracentered (meaning that a cell centred on the camera detector remains centred when you change objective). If it is not the case, you need to calibrate this difference and integrate it in the macro.*

#### 5.8. Pairing of the matching structures

First, the contours obtained in both modalities need to be paired. A first proposal is made by matching areas of the structures. The user can modify the correspondence (more details here).

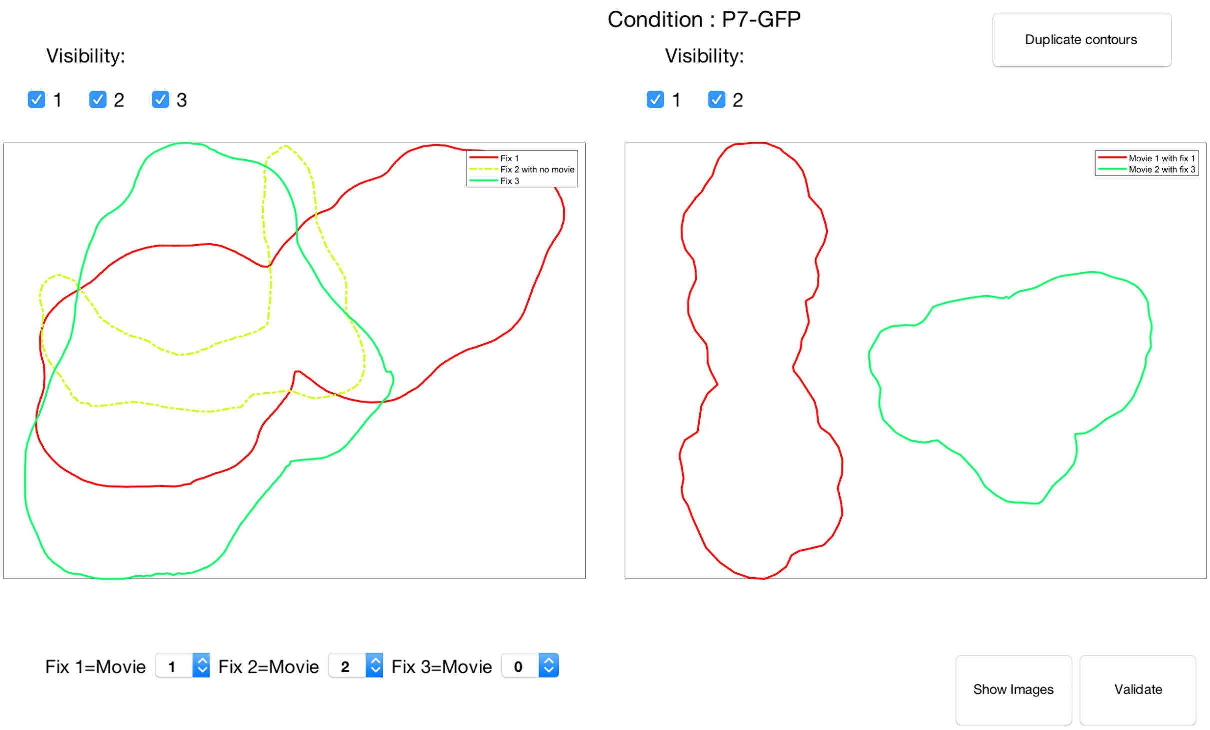

#### 5.9. Alignment of the paired structures

The matching contours then need to be aligned; an automatic method is coded but a manual alignment can also be performed, based on Flip/Translation/Rotation (more details [here]).

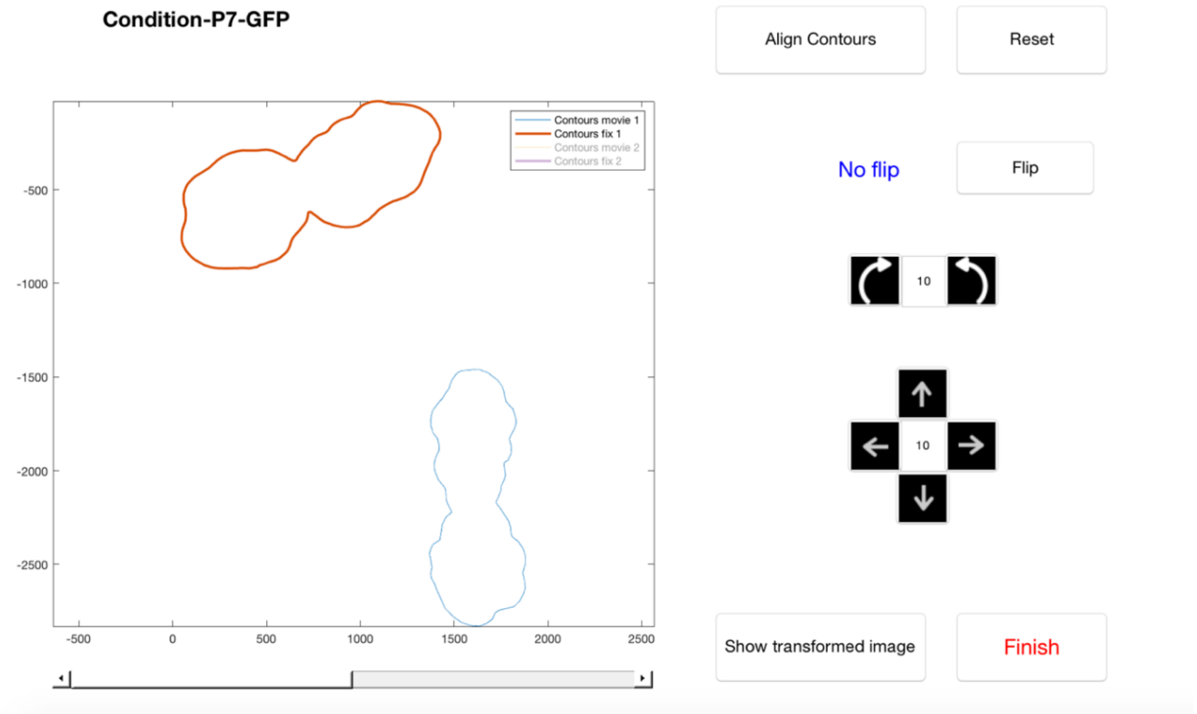

These two interfaces appear successively for each condition.

#### 5.10. Application of the transformation on images

In the ImageJ interface, the user choses which image should be aligned (to correspond to the other). Depending on this choice, the correct transformation is applied on the correct images. The results are saved in the folder *Results* in the main directory.

### 6. Detailed presentation of the three interfaces

#### 6.1. Presentation of the ImageJ interface

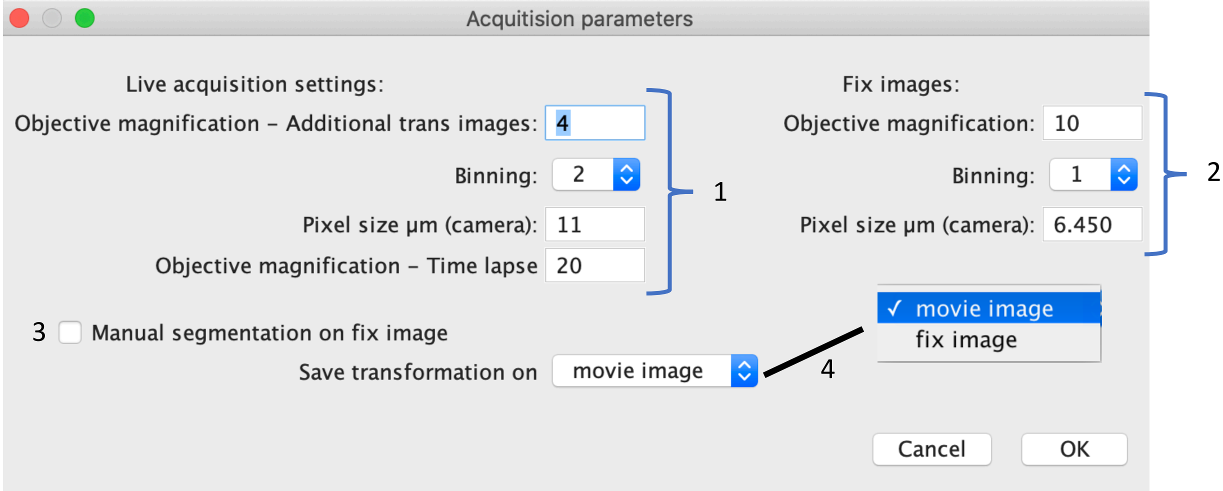

##### 6.1.1. <u>Information about the live acquisition & 2/ Information about fix images</u>

Are asked: the objectives used for each acquisition and some camera parameters (pixel size and binning):

- Objective magnifications determine the sub-folder names in the main directory
- These values enable the conversion pixel/µm (for the position stage, for computing the areas of structures, for the representation of the movie area on the reconstruction image)

##### 6.1.2. <u>Segmentation on the fix mosaic images</u>

The segmentation on the fix image is normally automatic but we also offer the possibility to do it manually. If checked, the macro will stop for each image in each sub-folder of *Directory_images_modality2* and ask the user to draw the contours. If a contour already exists, the macro loads it automatically.

##### 6.1.3. <u>Saving results</u>

Two options are proposed for the results:

- Rotate the multi-stage acquisition reconstruction (first modality): “movie image”
- Rotate the images of the second modality (mosaic-fix sample): “fix image”

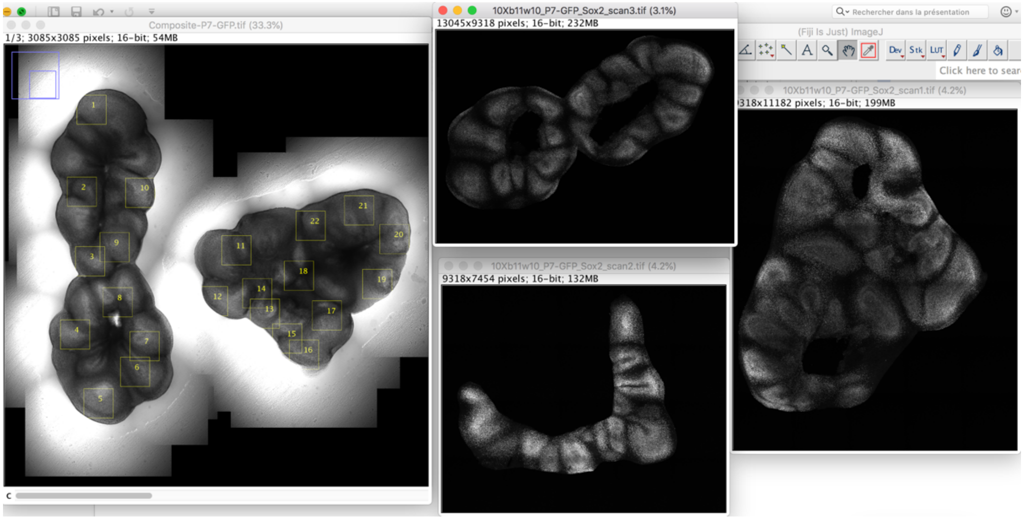

Example with 2 structures in the first acquisition image (2 tissue slices live-imaged) and 3 structures (3 tissue slices within the condition) in the second one.

If « *fix image* » is chosen, the two corresponding fluorescence images are rotated to correspond to the transmitted light images reconstruction:

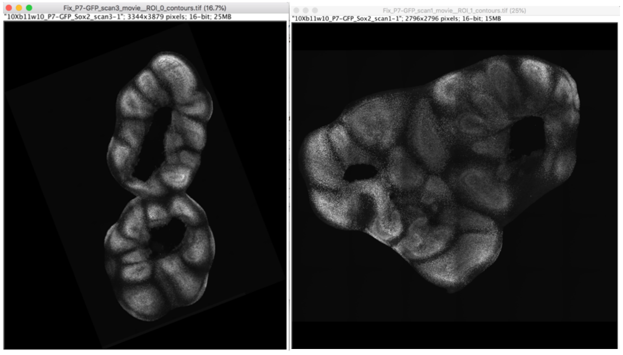

If «*movie image*» is chosen, the reconstruction of the first modality is rotated to correspond to each one of the mosaic fix structure (two rotations in this case because two structures in the reconstruction of the 1$^{st}$ modality). The squares highlight the targeted shape:

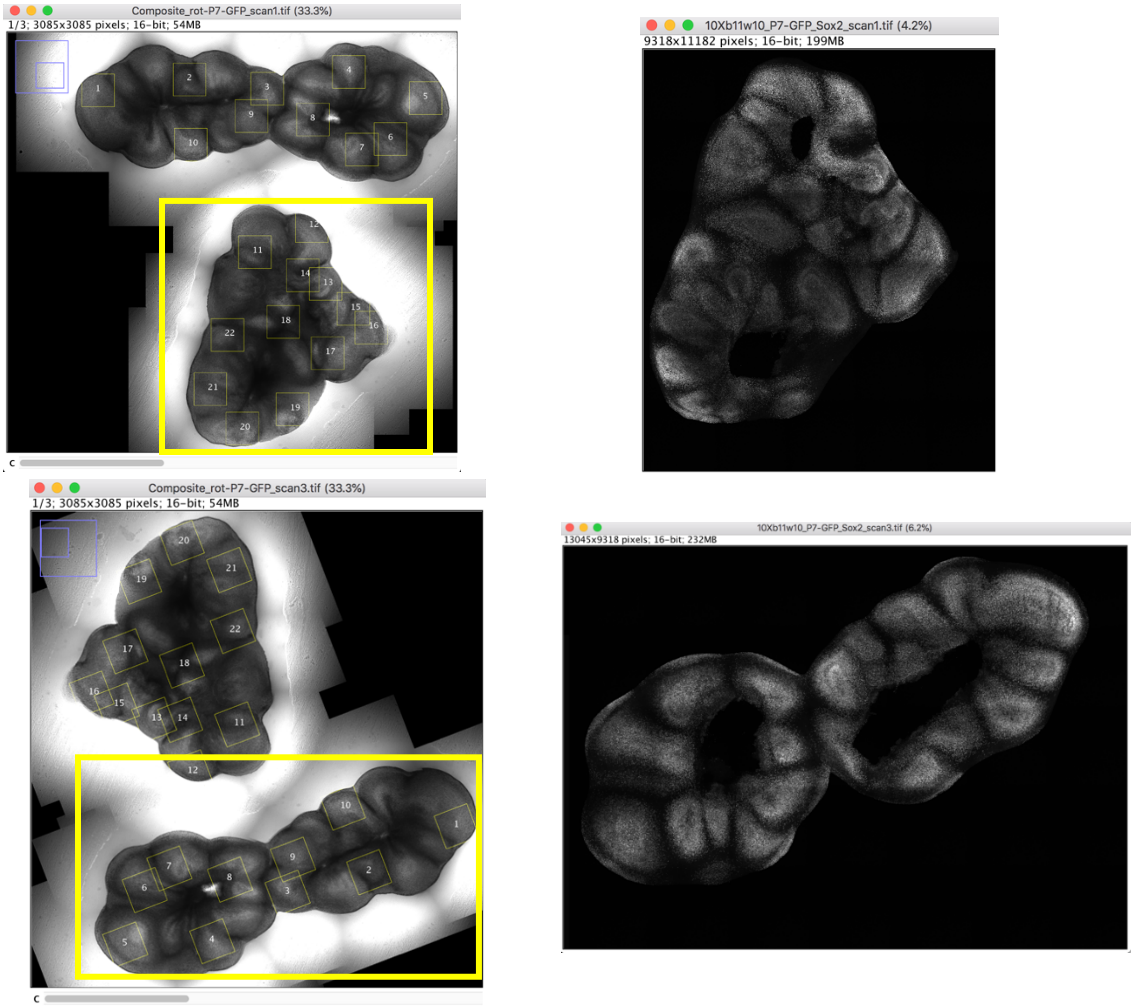

#### 6.2. Presentation of the first MATLAB interface: pairing of the contours

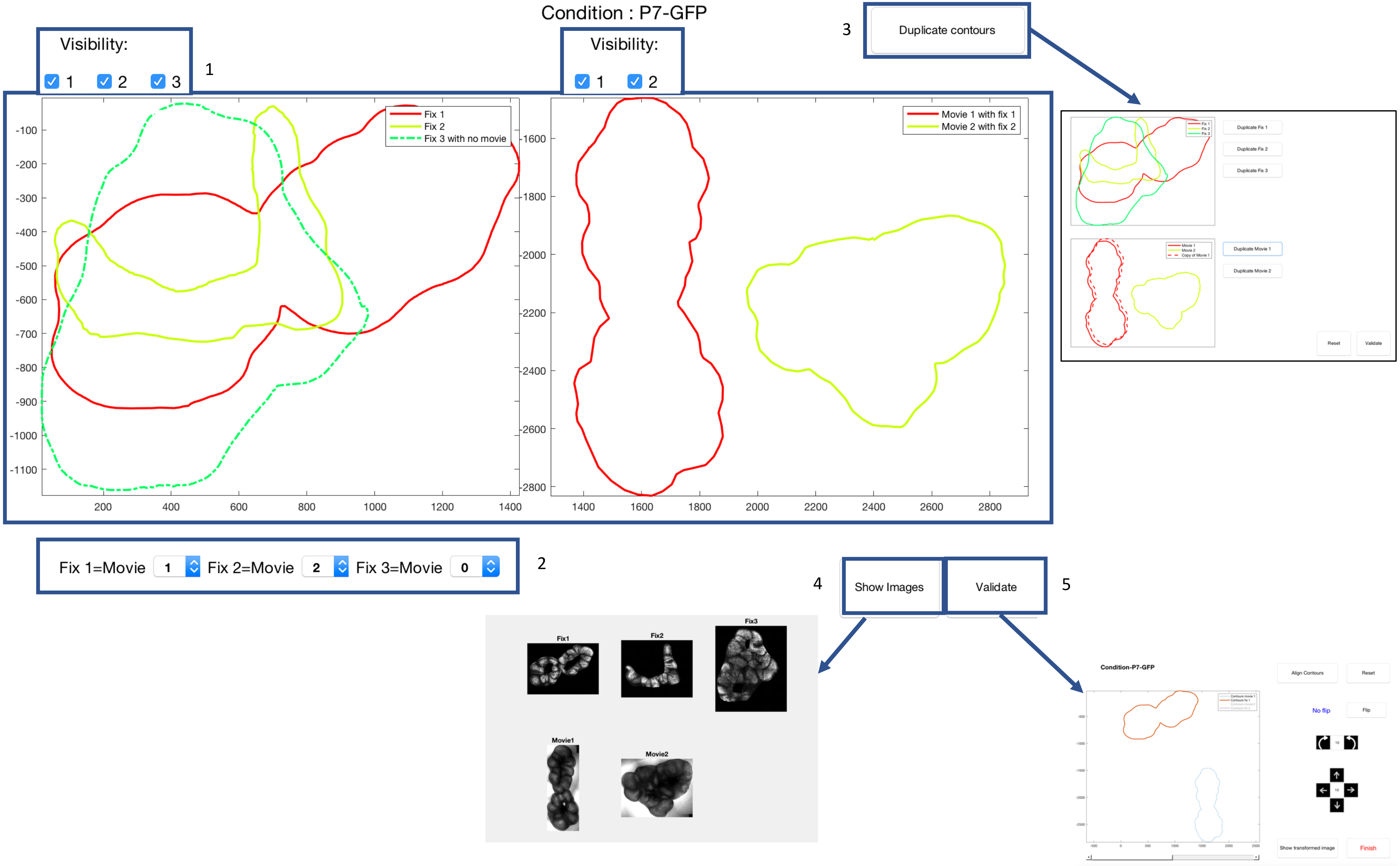

This interface enables to pair the contours from both modalities. A first proposal is made by the script minimizing:

$$\min_{i,j} (\mathcal{A}_1(i) - \mathcal{A}_2(j))^2,$$

where $\mathcal{A}_1$represent the areas corresponding to the first modality and $\mathcal{A}_2$ the areas corresponding to the second modality. The user can easily modify the association. Indeed, variations of the area were frequently observed between the first and the second modality. This phenomenon is caused by the mounting step of the immunostaining, where the coverslip slightly flattens the tissue. Thus, for the same tissue slice, the area of the second modality was often bigger than the area of the first modality.

###### 6.2.1. Display and visibility of contours

The contours are displayed in two different panels: one modality on one side, the other on the other side. When there is a match, the contours have the same color in both panels; otherwise, the contour is displayed in another color and with dotted lines. For a better visibility, the displayed contours on each panel can be selected by checking and unchecking the dedicated boxes "Visibility".

***Warning!*** *The total number of contours in each modality cannot exceed 9.*

###### 6.2.2. Association of the contours

The association is made on the contours of the second modality (the "fix" ones). The user can associate to each fix contour a movie contour (if no association: 0). Each contour can be paired only once.

###### 6.2.3. Duplicate contours

During the process of fixation and staining, the tissue slices can occasionally break or fold on themselves, making the pairing step impossible (since each contour can be used only once). We therefore gave the possibility to duplicate contours in order to treat fragmented tissue slices.

When clicking on "Duplicate contours", a new window opens with a button for each original contour, enabling to copy it. If clicked, the copy will be drawn in non-continuous points, with the same color, next to the original contour.
The "Reset" button reinitializes the interface: all duplicated contours are removed.
The "Validate" button adds the additional contour(s) to the original ones in the main window.
***Warning!*** *this change is definitive (but it is still possible to not associate this contour to a movie).*

Since the interface cannot treat more than 9 contours at the same time, if/when the number of contours in one of the modalities reaches 9, it is not possible to duplicate anymore.

###### 6.2.4. Show images

To help the user with the association (especially when several tissues have similar shapes), the images corresponding to the represented contours may be displayed in a new window thanks to the button "Show images".

###### 6.2.5. Validate

This button launches the second interface that enables to align the pairs of contours.

#### 6.3. Presentation of the second MATLAB interface: alignment

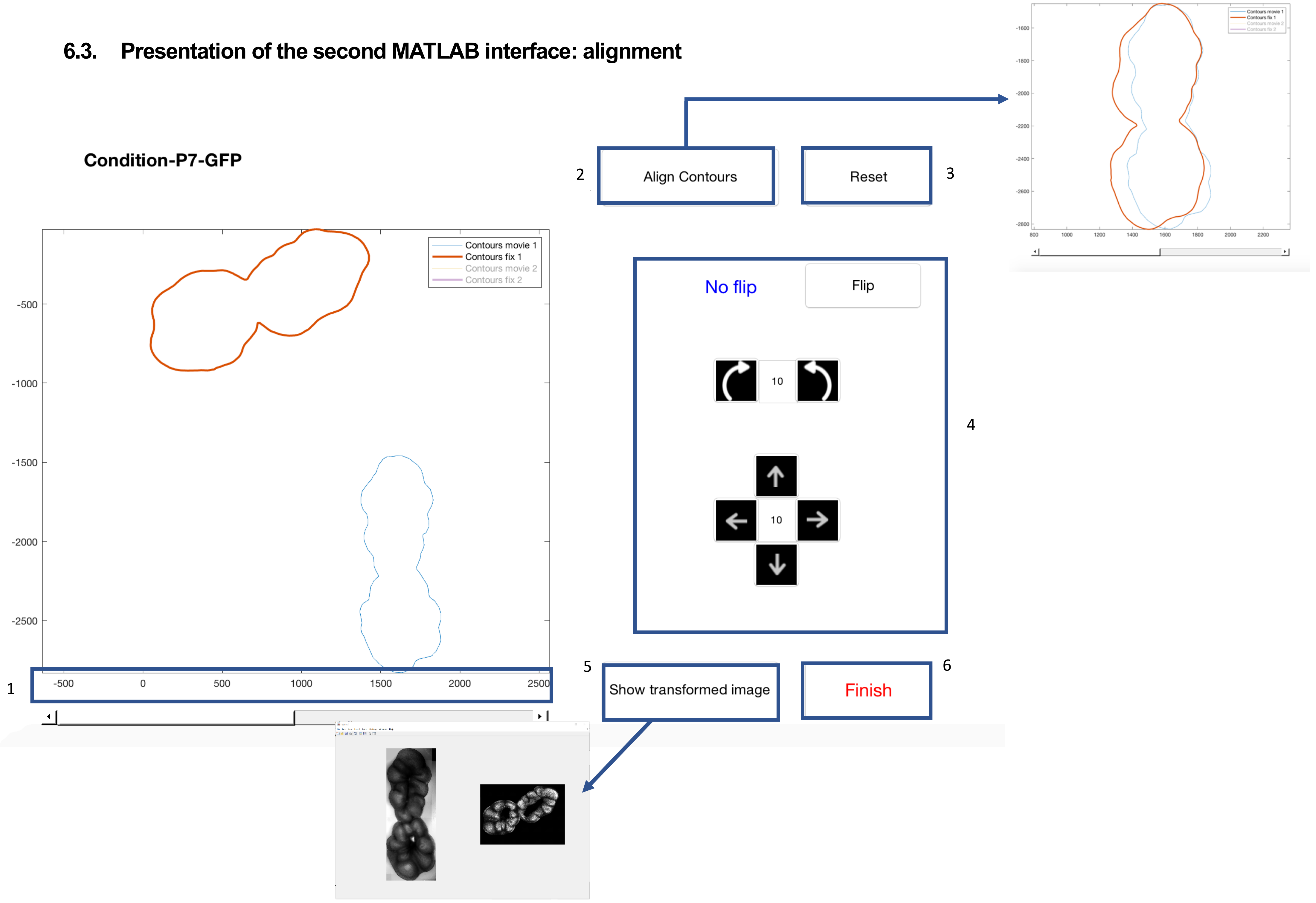

For each pair of contours associated in the previous interface, this interface proposes an automatic or manual alignment of these contours.

##### 6.3.1. Slider on image pairs

Enables to see the different pairs chosen in the previous interface. Each time you change the position of the slider, the displayed contours change. In this example from the previous interface, we had two pairs: the slider therefore has two positions.

***Warning!*** *Each pair of contours must be treated to close this interface.*

##### 6.3.2. Align contours button

Automatically aligns the displayed contours (details on the algorithm here). This automatic alignment can be done only once for each pair of contours; the button is afterwards “disabled”. Manual adjustments can be performed on the displayed result.

##### 6.3.3. Reset button

Resets the transformation (for the displayed pair of contours) and shows the contours in their initial state.

##### 6.3.4. Manual alignment panel

Offers the possibility to align manually the contours, based on 3 transformations: translation, rotation and flip. The translation step and the rotation angle can be changed in the dedicated boxes.

##### 6.3.5. Show transformed image button

Shows, for the displayed pair of contours, the corresponding images: on the left, the reference image (in our case the transmitted light acquisition) and on the right, the other image (in our case, the fluorescence image) **after transformation** so that the contour matches the representation in the interface. It always corresponds to the contour displayed in the main interface at the time the button is clicked (no live update).

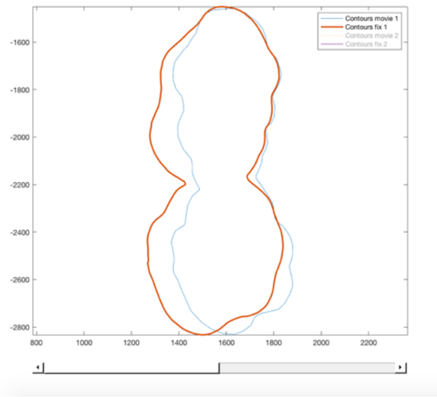

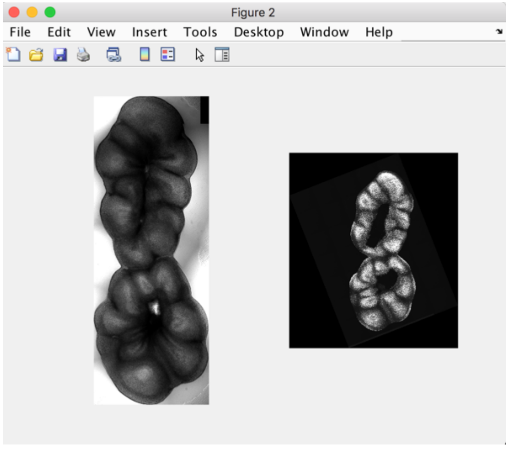

##### 6.3.6. Finish button

Once satisfied with the alignment on **all pairs** of contours, click *Finish*; if there is another condition to treat, the first MATLAB interface will appear with the contours corresponding to the next condition. Otherwise this part is over and the ImageJ macro finishes the process.

If no transformation has been applied to one of the pairs, you will have an error message:

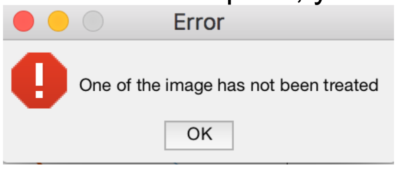

One cannot finish the alignment process if no transformation has been applied to one pair of contours.

#### 7. Algorithm details on contours alignment

According to the biological model, between the two modalities, the possible mathematical operations are: translation, rotation and flip; because the scale can be a bit different, we also consider scaling transformation. We used the MATLAB function “*fitgeotrans*” computing the best geometrical transformation between two **ordered** sets of points. According to our hypothesis, we chose the ‘similarity’ transformation. Arbitrarily, the first modality contour is considered as the “reference” one and the second modality contour is transformed to be as close as possible to the reference one.

The “*fitgeotrans*” function computes the best transformation to apply to a set of points to correspond to another set of points, matching points 2 by 2, in the given order (first points of each set together, second points together etc.):

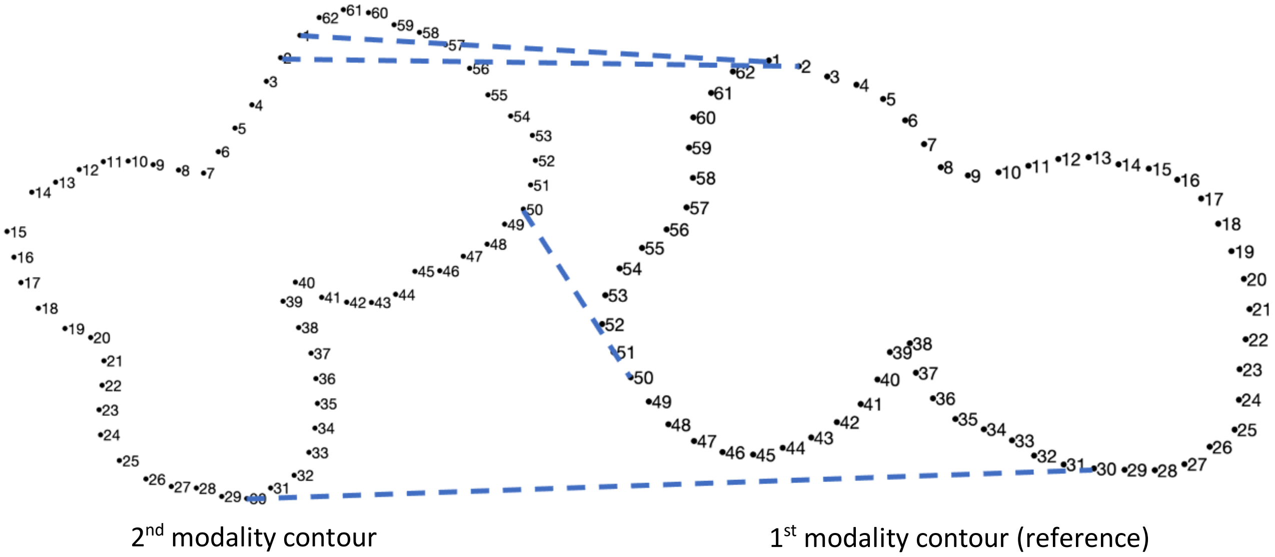

However, the contours coming from the macro may not have the same starting points (and/or order) and even the same number of points. A first pre-processing is therefore applied on both contours so that they have the same number of points, evenly distributed.

To find the matching first points, all possible (circular) translations of the set of points are created (the second original point becomes the first one, then the third original one becomes the first one etc., until the end of the original contour):

Original contour : (X_f (1),Y_f (1)),(X_f (2),Y_f (2)),… ,(X_f (n-1),Y_f (n-1)),(X_f (n),Y_f (n))
First translation : (X_f (2),Y_f (2)),(X_f (3),Y_f (3)),… ,(X_f (n),Y_f (n)),(X_f (1),Y_f (1))
…
Last translation : (X_f (n),Y_f (n)),(X_f (1),Y_f (1)),… ,(X_f (n-2),Y_f (n-2)),(X_f (n-1),Y_f (n-1))

The same computation is performed on the “inverse” (flipped) set of points (going in the other direction, see representation below):

First inversion : (X_f (n),Y_f (n)),(X_f (n-1),Y_f (n-1)),… ,(X_f (2),Y_f (2)),(X_f (1),Y_f (1))
Second inversion : (X_f (n-1),Y_f (n-1)),(X_f (n-2),Y_f (n-2)),… ,(X_f (1),Y_f (1)),(X_f (n),Y_f (n))
…
Last inversion : (X_f (1),Y_f (1)),(X_f (n),Y_f (n)),… ,(X_f (3),Y_f (3)),(X_f (2),Y_f (2))

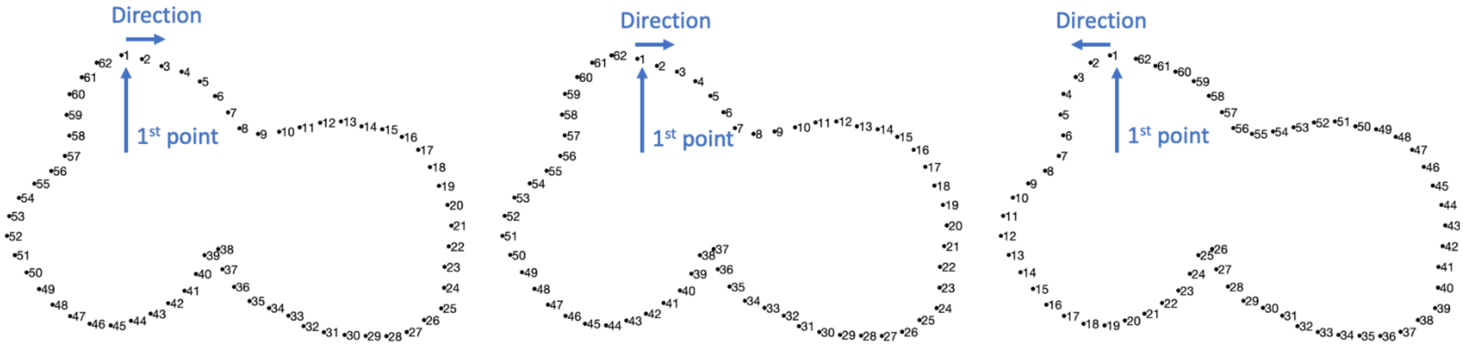

Original contour (1 point over 20) 1st translation Last contour inversion

To determine the best combination, an energy E is associated to each one, based on the Euclidian distance between the reference set of points (the points describing the contour of the first modality, $(X_1, Y_1)$) and the transformed set of points of the second modality, $(X_2, Y_2)$, after transformation $t$.

$$\min_{t} E\big((X_2, Y_2), (X_1, Y_1), t\big),$$

$$\text{with } E\big((X_2, Y_2), (X_1, Y_1), t\big) = \sqrt{\sum_i \ (X_1(i) - t(X_2)(i))^2 + (Y_1(i) - t(Y_2)(i))^2}\ ,$$

where $t$ represents the transformation applied to the set of points, composed of:

- Circular translation and inversion
- Transformation computed by *fitgeotrans.*

For the example above, the combination with the lowest energy is shown on the left (original contour was inversed and translated) and the contour after transformation on the right:

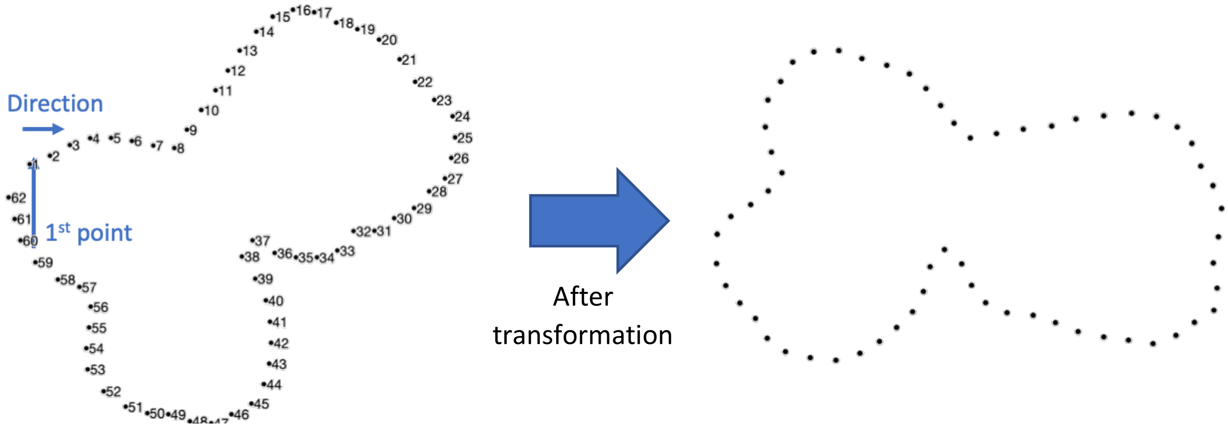

Ordered set of points minimizing the energy 2nd modality contour after transformation

The images are then correctly aligned:

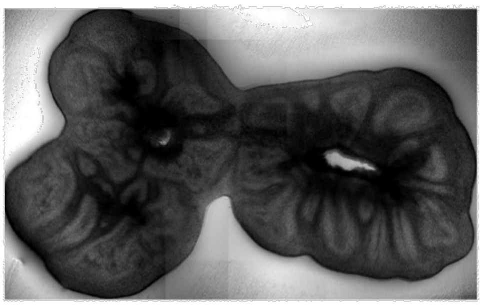

1st modality contour (reference)

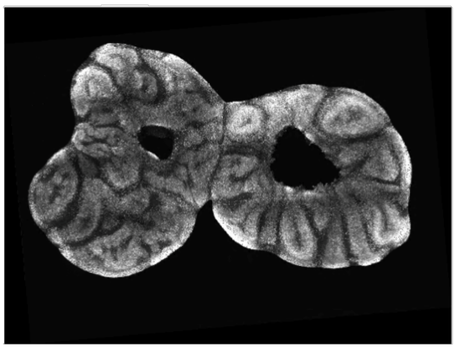

2nd modality contour after transformation

The transformation is applied to the original images in the ImageJ macro. The rotation and potential flip must be returned. The transformation matrix *R* between the two contours can take two forms:

$$\text{No flip: } R = \alpha\begin{pmatrix} \cos(\theta) & \sin(\theta) & t_x \\ -\sin(\theta) & \cos(\theta) & t_y \\ 0 & 0 & 1 \end{pmatrix} \qquad \text{flip: } R = \alpha\begin{pmatrix} \cos(\theta) & \sin(\theta) & t_x \\ \sin(\theta) & -\cos(\theta) & t_y \\ 0 & 0 & 1 \end{pmatrix}.$$

The angle of rotation $\theta$ is given by the formula (the scaling factor is not considered):

$$\theta = \arctan\left({}^{R(1,2)}\big/_{R(1,1)}\right).$$

For each condition, a text file read by the ImageJ macro is created by MATLAB giving the potential flip (1 or 0) and the rotation angle:

| | Name | Value |
|---|---|---|
| 1 | nbImages | 3 |
| 2 | Images1_path | {...}\10Xb12w9_P2-MSCV-GFP_Sox2_scan1.tif |
| 3 | Images1_rotAngle | -8 |
| 4 | Images1_Flip | 1 |
| 5 | Images2_path | {...}\10Xb12w9_P2-MSCV-GFP_Sox2_scan2.tif |
| 6 | Images2_rotAngle | 3 |
| 7 | Images2_Flip | 1 |
| 8 | Images3_path | {...}\10Xb12w9_P2-MSCV-GFP_Sox2_scan3.tif |
| 9 | Images3_rotAngle | -61 |
| 10 | Images3_Flip | 0 |

*Remark: This file is read by the macro to apply the transformations on the images (step 5.10).*

### 8. Easy changes in the code

The ImageJ macro was written so that some functionalities can be easily modified to use it for other applications, with the same purpose: finding transformation between pairs of contours (representing the same structures) acquired in different modalities. Even if our code was created for transmitted light microscopy and fluorescence microscopy, we offer a flexible tool that could be used with other type of images.

For any questions or adaptation about this code, please feel free to contact us: ** or **

#### 8.1. About the second modality acquisition

- **How to change the (automatic) segmentation?**

The segmentation on these images was adapted to our acquisition (the steps were explained in paragraph 5.6).
Function to modify: segmentFluoImages
Requirements: the contours (set of coordinates) are saved in the subfolder *Results* in the folder containing the images, as X and Y coordinates

- **How to use it with images acquired with different modality?**

Choose the "manual segmentation" and this should work perfectly; if the segmentation can be automatic, do not hesitate to change the function as described just before.

- **How to remove/change the subsampling applied on these images?**

Because in our case the images were quite big (mosaic acquisition), we subsampled the images by a factor 4, simply to decrease the computational time.
Parameter to modify: "*p_sech*" (line 10) at the beginning of the code which corresponds to the inverse of the value you would enter in Image->Scale; put 1 if you want no subsampling.

#### 8.2. About the STG file

- **Use a file of positions different of a "STG" file as described here?**

We made some tests using *NIS.Elements* software. In this case, the generated file is a XML file with a lot of parameters (not only the stage positions) and we decided to make a small script in MATLAB to generate a STG file so that the macro remains usable exactly the same way.

- **Change pattern for condition name recognition in the STG file?**

The variable "*str_afterCondName_STG*" indicates which strings delimit the name of the condition read in the STG file (the condition name is the string found before this delimiter):
Parameter to modify: "*str_afterCondName_STG*" (line 12)
Be careful, this string must be present in each position name.

#### 8.3. About the first modality acquisition

- **Change the (manual) segmentation?**

Function to modify: drawContoursTransImage
Requirements: that the contours are saved in the subfolder *Results* in the folder containing the images, as X and Y coordinates.

- **Change the type of file for the multi-position images?**

The data of the first modality are loaded using the nd file of the acquisition.
Function to modify: openTransAcquisition_oneCondition
Requirements: the images should be organized as a stack image at the end of the function (because a z-projection is applied afterwards).
